# Wasabi leaf extract changes the expression levels of cytokine related genes in dermal papilla cells, which identified by the whole transcriptome

**DOI:** 10.1101/2025.05.20.653605

**Authors:** Himari Matsusaka, Tomoe Yamada-Kato, Mayuko Nakai, Lanlan Bai, Hiroshi Tomita, Eriko Sugano, Taku Ozaki, Isao Okunishi, Tomokazu Fukuda

## Abstract

Wasabi is a traditional food in Asian area. Wasabi leaf extract derived compounds such as 6-methylsulfinylhexyl isothiocyanate (6-MSITC) have been reported to exert anti-proliferative effects on cancer cells. The antibacterial action of wasabi leaf extract has been reported previously. However, the detailed biological effects of wasabi leaf extract are not fully understood. To get a clue for the understanding, we used next-generation sequencing to identify the upregulated or downregulated genes in immortalized human dermal papilla cells. Differentially expressed genes were identified in human papilla cells in the presence or absence of wasabi leaf extract. The effects of 6-MSITC and isosaponarin were evaluated. We found that wasabi leaf extracts upregulated or downregulated the expression of cytokine-related genes. The identified expression changes will contribute to our understanding of the biological effects of wasabi.

## INTRODUCTION

The wasabi is a native and traditional plant of Asian countries. The cultivation of wasabi started around 16-17^th^ century (Shizuoka, 2023). The strong flavor of wasabi is internationally recognized as an essential compliment to sushi (Morgan, 2005). Therefore, understanding the flavor of wasabi is essential to understand the cultural background of traditional Asian foods.

The biological effects of wasabi have been studied in various cell culture experiments. In brief, 6-methylsulfinylhexyl isothiocyanate (6-MSITC), a major component of wasabi root and rhizome, has been reported to have anti-proliferative effects against gastric cancer-derived cells (Watanabe et al., 2003). Furthermore, antioxidant effects of 6-MSITC via the Nrf2-mediated pathway have been reported in HepG2 cancer-derived cells (Trio et al., 2017). This information can help understand the biological effects of wasabi root and leaf extracts. However, to the best of our knowledge, no study has described the effect of wasabi leaf extracts from view point of transcriptome based on the next-generation sequencing.

Our research group previously reported that expression of human derived R24C mutant expression of cycline dependent kinase 4 (CDK4), cyclin D1, and telomerase reverse transcriptase (TERT) allows efficient immortalization with various types of cells including myogenic (Shiomi et al., 2011), corneal epithermal (Furuya et al., 2022), dental pulp (Orimoto et al., 2020), airway epithermal (Orimoto et al., 2021), and dermal papilla cells (Fukuda et al., 2020). The immortalization of the expression of these three genes was named K4DT based on the last characters of the gene name. Although the expression of traditional virus-derived oncogenes, such as SV40 or E6E7, frequently cause chromosomal abnormalities, the expression of K4DT showed that chromosomal conditions are relatively intact (Fukuda et al., 2021). Furthermore, we successfully established the dermal papilla cell that constitutively expressed the androgen receptor (AR) using the retroviral gene transfer method. Exposure of dihydroxy testosterone (DHT) causes efficient translocation into the cytoplasm and nucleus, suggesting that our introduced AR works properly as a steroid receptor (Fukuda et al., 2020). In this study, we exposed wasabi leaf extract and performed RNA-Seq analysis to identify the biological effects of wasabi leaf extract.

## MATERIALS AND METHODS

### Cell lines and treatments

Original primary cells were purchased from PromoCell (Heidelberg, Germany) using a local distributor (Takara Bio, Shiga, Japan). Detailed information on immortalized human dermal papilla cells (DPCs) expressing mutant CDK4, Cyclin D1, and TERT has been previously reported (Fukuda et al., 2020). Furthermore, we established immortalized DPCs expressing the AR via retroviral gene transfer (Fukuda et al., 2020). We established 15 experimental samples to cover all aspects of the biological response to AR signal activation. To ensure reproducibility, each experimental group consisted of three samples.

Sample 1: HFDPC_K4DT_AR_rep1, AR, immortalized DPCs with retroviral AR expression, no ligand treatment.
Sample 2: HFDPC_K4DT_AR_rep2, AR, immortalized DPCs with retroviral AR expression, no ligand treatment.
Sample 3: HFDPC_K4DT_AR_rep3, AR, immortalized DPCs with retroviral AR expression, no ligand treatment.
Sample 4: HFDPC_K4DT_AR_DHT_rep1, ARDHT, immortalized DPCs with retroviral AR expression treated with 50 nM DHT.
Sample 5: HFDPC_K4DT_AR_DHT_rep2, ARDHT, immortalized DPCs with retroviral AR expression treated with 50 nM DHT.
Sample 6: HFDPC_K4DT_AR_DHT_rep3, ARDHT, immortalized DPCs with retroviral AR expression treated with 50 nM DHT.
Sample 7: HFDPC_K4DT_AR_DHT_iso_rep1, ARDHT, immortalized DPCs with retroviral AR expression, treated with 50 nM DHT and 10 μM isosaponarin.
Sample 8: HFDPC_K4DT_AR_DHT_iso_rep2, ARDHT, immortalized DPCs with retroviral AR expression, treated with 50 nM DHT and 10 μM isosaponarin.
Sample 9: HFDPC_K4DT_AR_DHT_iso_rep3, ARDHT, immortalized DPCs with retroviral AR expression, treated with 50 nM DHT and 10 μM isosaponarin.
Sample 10: HFDPC_K4DT_AR_DHT_6-MSITC_rep1, ARDHT, immortalized DPCs with retroviral AR expression, treated with 50 nM DHT and 50 nM 6-MSITC (6-methylsulfinyl hexyl isothiocyanate).
Sample 11: HFDPC_K4DT_AR_DHT_6-MSITC_rep2, ARDHT, immortalized DPCs with retroviral AR expression, treated with 50 nM DHT and 50 nM 6-MSITC (6-Methylsulfinyl hexyl isothiocyanate).
Sample 12: HFDPC_K4DT_AR_DHT_6-MSITC_rep3, ARDHT, immortalized DPCs with retroviral AR expression, treated with 50 nM DHT and 50 nM 6-MSITC (6-Methylsulfinyl hexyl isothiocyanate).
Sample 13: HFDPC_K4DT_AR_DHT_wasabi_rep1, ARDHT, immortalized DPCs with retroviral AR expression, treated with 50 nM DHT and 0.06% wasabi leaf extract.
Sample 14: HFDPC_K4DT_AR_DHT_wasabi_rep2, ARDHT, immortalized DPCs with retroviral AR expression, treated with 50 nM DHT and 0.06% wasabi leaf extract.
Sample 15: HFDPC_K4DT_AR_DHT_wasabi_rep3, ARDHT, immortalized DPCs with retroviral AR expression, treated with 50 nM DHT and 0.06% wasabi leaf extract.

The expression profiles of Sample 1-6 were previously obtained and reported by our group as an independent manuscript that focused on the androgen receptor signaling pathway (Matsusaka et al., 2022). In this study, we compared the expression profiles of isosaponarin, 6-MSITC, and wasabi leaf extract.

### Treatments of cells and RNA extraction

Cells were treated with DHT or DHT plus isosaponarin, DHT plus 6-MSITC, or DHT plus wasabi leaf extract for 24 h at approximately 70% confluence in a 35 mm diameter cell culture dish. The cells were then treated with 50 nM DHT. Isosaponarin (10 μM) and 6-MSITC (50nM) treatments were conducted in addition to the DHT treatment. Total cellular RNA was extracted using the Nucleopsin RNA kit (Takara Bio) according to the manufacturer’s instructions. DNase I treatment was performed using enzymes included in the kit. Total RNA quality was assessed using tapestation and cDNA libraries were prepared using the NEBNext Ultra II Directional RNA Library Prep Kit for Illumina (New England BioLabs, Ipswich, Massachusetts, USA). The RIN value of the RNA extracted from all samples was 10.0.

### Sequencing reaction and processing of the data

A cDNA library was created using poly A-tailed primers. The HiseqX sequencer (Illumina, San Diego, CA, USA) was used to obtain pair-end 150 bp sequencing reads. Output reads were processed using the PEAT program to remove the adaptor sequences (Li et al., 2015). Low-quality reads were filtered using FASTP (Chen et al., 2018). The filtered reads were uploaded to the NCBI SRA database using the BioProject PRJNA1005423 and Submission SUB13751027. We have prepared a link (https://dataview.ncbi.nlm.nih.gov/object/PRJNA1005423?reviewer=3vult744ghtvpaq2m0qfue62ll). Filtered reads were mapped to the NCBI reference human genome sequence (CRCh38) using STAR (Dobin et al., 2013). The mapped data were processed into expression count data using the feature *Counts* in the subread package (Liao et al., 2014). To extract differentially expressed genes (DE genes), we performed a pairwise comparison between HFDPC_K4DT_AR_DHT, ARDHT, and immortalized DPCs with retroviral AR expression and HFDPC_K4DT_AR_DHT, ARDHT, and immortalized DPCs with retroviral AR expression treated with 50 nM DHT and 10 μM isosaponarin.

Furthermore, we compared two samples: HFDPC_K4DT_AR_DHT, ARDHT, and immortalized DPCs with retroviral AR expression and HFDPC_K4DT_AR_DHT_rep1, ARDHT, and immortalized DPCs with retroviral AR expression treated with 50 nM DHT and 50 nM 6-MSITC. Finally, we compared HFDPC_K4DT_AR_DHT, ARDHT, and immortalized DPCs with retroviral AR expression and HFDPC_K4DT_AR_DHT, ARDHT, and immortalized DPCs with retroviral AR expression treated with 50 nM DHT and 0.06% wasabi leaf extract. iDep (http://bioinformatics.sdstate.edu/idep96/) was used for comparison. FDR threshold was set to 0.05. We also processed the data using TCC-GUI for principal component analysis (PCA), heatmap analysis, and bar plot analysis (Su et al., 2019).

### Downstream analysis

After the identification of DE genes, we applied the upregulated or downregulated genes for functional annotation. The Database of Functional Annotations, DAVID, was used for annotation. Based on the pathways listed in DAVID, we marked the pathway positions with arrows in the corresponding Figures.

### Chemicals and reagents

Isovitexin (6-C-glucosyl apigenin) was purchased from Extrasynthese (Lyon, France). Isoorientin (6-C-glucosyl luteolin) was obtained commercially (Toronto Research Chemicals, Toronto, Canada). Isosaponarin (4’-O-glucosyl-6-C-glucosyl pigenin) was purified from wasabi leaves according to a previously described method (Nagai et al., 2010). We determined the purity of isosaponarin and 6-MSITC with nuclear magnetic resonance spectroscopy.

### Preparation of wasabi leaf extract

Wasabi was cultivated and harvested in the spring of 2019 in Mie, Japan. The crushed leaves were mixed with hot water at a ratio of 2:5 and extracted above 85 ℃ for 10 min. After filtration through a filter cloth, the extracted liquid was concentrated to approximately 1/5 of the leaves. For sterilization and precipitation, the concentrated liquid was autoclaved at 123 ℃ for 60 min. After filtration through a 1 μm filter, the liquid was freeze-dried and powdered. The Wasabi leaf extract was frozen until the start of the study.

### Preparation of sample solution for Liquid Chromatography-Mass spectrometry (LC/MS)

The wasabi leaf extract (0.1 g) was ultrasonically extracted in 20 mL of water. The sample solution was prepared by filtration using a 0.22 μm hydrophilic polytetrafluoroethylene (PTFE) syringe filter (GLCTD-PTFE1322, Shimazu GLC Ltd., Tokyo, Japan).

### LC/MS analysis

Chromatographic analysis was performed using a Thermo Fisher Scientific UltiMate 3000 Rapid Separation LC System (Thermo Fisher, MA, USA). The separation was carried out on a Develosil C30-UG-5 column (250 x 4.6 mm ID; Nomura Chem. Co., Ltd., Aichi, Japan). The mobile phase was composed of ultrapure water (0.1% formic acid, A) and methanol (0.1% formic acid, B), using a gradient elution of 2% B at 0–2.5 min, 70% B at 62 min, and 98% B at 68–80 min. The sample injection volume was 1 µL. The UV signal was measured at 254 nm. An ESI-Q-ToF Maxis (Bruker Daltonics, MA, USA) was connected to the LC system via an electrospray ionization (ESI) interface. The acquisition parameters were as follows: drying gas flow rate, 8.0 L/min; drying gas temperature, 200 ℃; nebulizer, 2.5 bar; capillary voltage, 4.5 kV. Data analysis was performed using DataAnalysis version 4.2 (Bruker Daltonics, MA, USA).

## RESULTS AND DISCUSSION

### The whole transcriptomes of AR expressing human dermal papilla cells

We subjected our cDNA to next-generation sequencing-based RNA-Seq. As shown in Figure 1A, the number of reads was at least 15M, which was sufficient for the quantitation of gene expression. The read quality data are presented in Figure S1–S3. The lowest mapping ratio was 95.2%, suggesting that the mapping process did not involve technical problems. As shown in Figure 1B, we compared the reproducibility of the replicates using matrix analysis. Three replicates formed a unique cluster in the analysis, indicating that the biological replication was reproducible within the experimental group. As shown in Figure 1C, PCA showed two major groups within the samples. The first group comprised of a sample without DHT treatment (HFDPC_K4DT_AR). The second group comprised of samples treated with DHT (HFDPC_K4DT_AR_DHT, HFDPC_K4DT_AR_DHT_iso, HFDPC_K4DT_AR_DHT_6-MSITC, and HFDPC_K4DT_AR_DHT_wasabi). The detailed location of the experimental groups in three dimensional visualization has been presented as a video (Movie1, Figshare, https://doi.org/10.6084/m9.figshare.25018175). Among the DHT-treated groups, the wasabi leaf extract-treated group was relatively distant from the other DHT-treated groups, such as HFDPC_K4DT_AR_DHT, HFDPC_K4DT_AR_DHT_iso, and HFDPC_K4DT_AR_DHT_6-MSITC.

**Figure 1.**
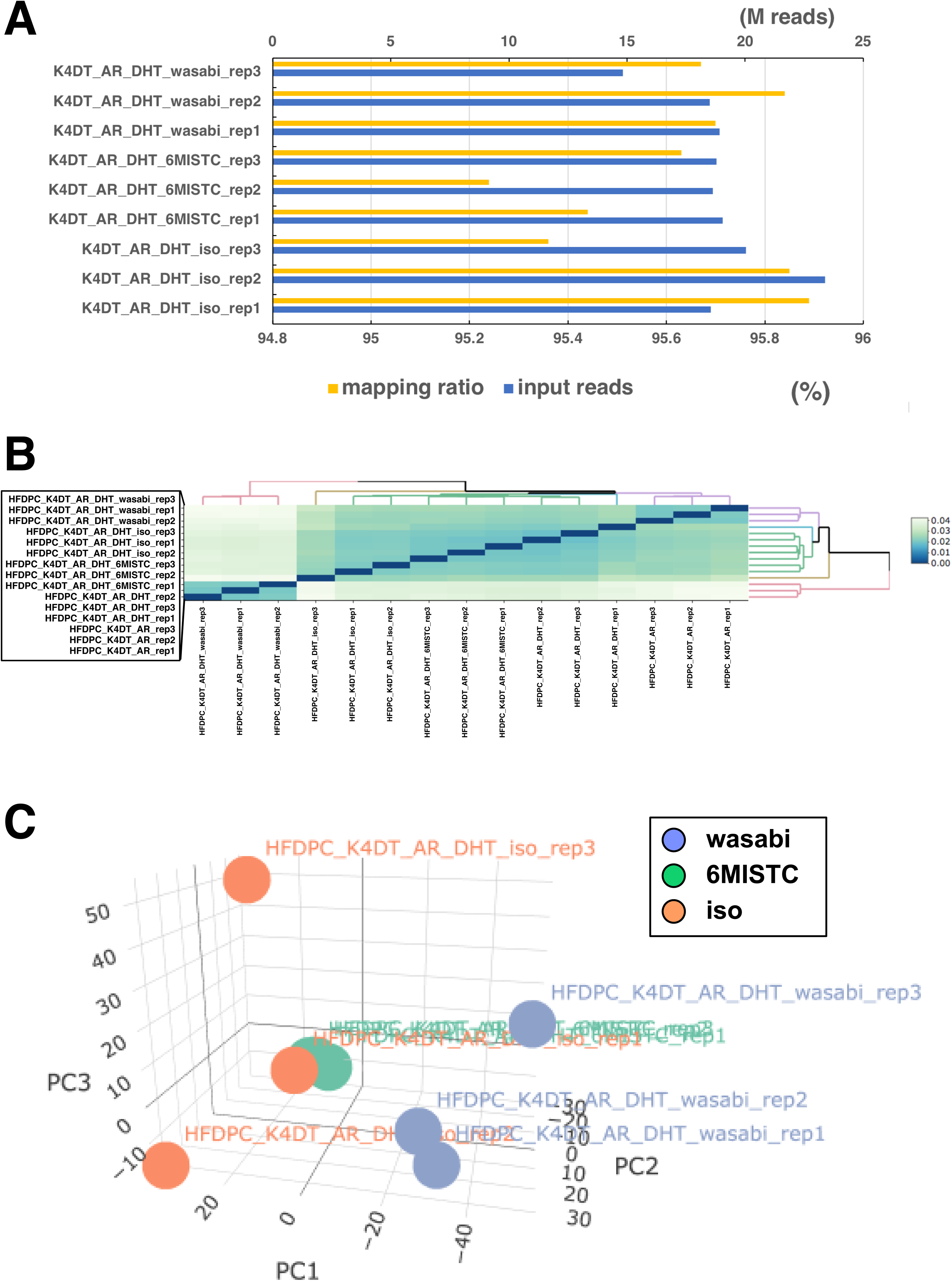
Mapping ratio, reproducibility and results of Principle Component Analysis (PCA) of the human derived immortalized dermal papilla cells (DPCs) on the transcriptome. A, Mapping ratio to target genome and number of input reads of each samples. B, Matrix analysis of all sample reads. C, PCA analysis of DPCs expressing AR (AR), DHT treated DPCs expressing AR (AR_DHT), DHT treated DPCs expressing AR under the existence of isosaponarin (iso), DHT treated DPCs expressing AR under the existence of 6-MSITC (6-MSITC), DHT treated DPCs expressing AR under the existence of wasabi leaf extract (wasabi).

### Extraction of differentially expressed genes in wasabi leaf extract treated group, isosaponarin treated group, 6-MSITC treated group

To identify the differentially expressed genes (DE genes) in the wasabi leaf extract-treated group, we performed pairwise comparisons of gene expression between HFDPC_K4DT_AR_DHT and HFDPC_K4DT_AR_DHT_wasabi using DE-Seq2 and iDep. As shown in Figure 2A, 487 upregulated (red bars) and 343 downregulated genes (blue bars) were identified at a significance level of 0.05. The number of DE genes was obtained from pairwise comparisons of HFDPC_K4DT_AR_DHT and HFDPC_K4DT_AR_DHT_wasabi, HFDPC_K4DT_AR_DHT, HFDPC_K4DT_AR_DHT_iso, HFDPC_K4DT_AR_DHT, and HFDPC_K4DT_AR_DHT_6-MSITC, as shown in Figure 2B. We observed that the number of DE genes was the highest in the comparison between HFDPC_K4DT_AR_DHT and HFDPC_K4DT_AR_DHT_wasabi, which is consistent with the results of the PCA.

**Figure 2.**
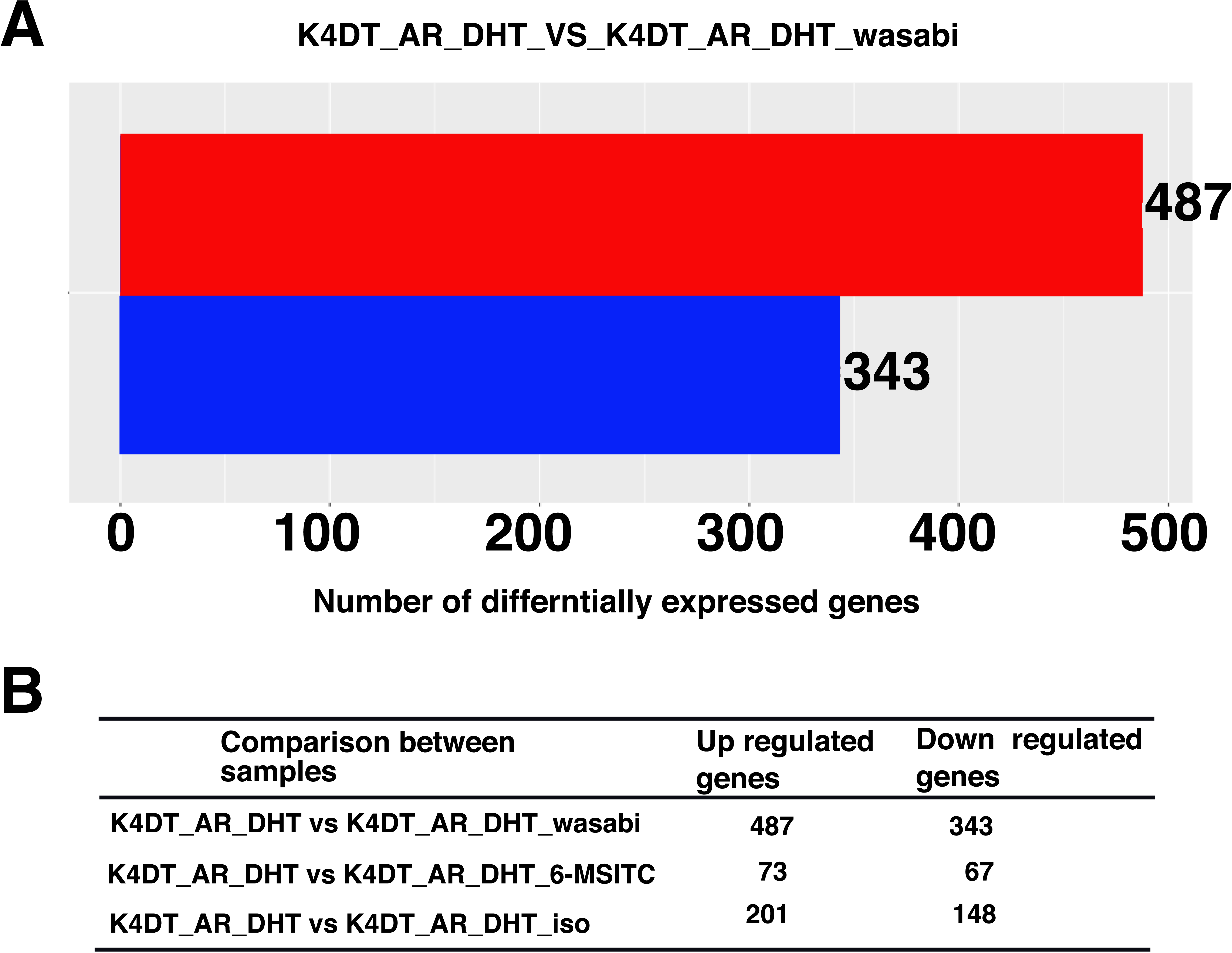
Results of pairwise comparison between DHT treated DPCs expressing AR (K4DT_AR_DHT) and DHT treated DPCs expressing AR under the exitance of wasabi leaf extract (K4DT_AR_DHT_wasabi). A, Number of differentially expressed genes (DE genes) on the pairwise comparison. The number of significant genes were counted. The red bar indicates the upregulated genes in wasabi treated group. The blue bar indicates the downregulated genes in wasabi treated group. B, The results of pairwise comparison between DHT treated DPCs expressing AR (K4DT_AR_DHT) and DHT treated DPCs expressing AR under the exitance of wasabi leaf extract (K4DT_AR_DHT_wasabi), between DHT treated DPCs expressing AR (K4DT_AR_DHT) and DHT treated DPCs expressing AR under the exitance of isosaponarin (K4DT_AR_DHT_iso), between DHT treated DPCs expressing AR (K4DT_AR_DHT) and DHT treated DPCs expressing AR under the exitance of 6-MSITC (K4DT_AR_DHT_6-MSITC).

### The heatmap analysis of DE genes in wasabi leaf extract treated group, isosaponarin treated group, 6-MSITC treated group

In Figure 3, we list the expression profiles of the 487 genes extracted from the pairwise comparisons between HFDPC_K4DT_AR_DHT and HFDPC_K4DT_AR_DHT_wasabi. The expression patterns in all experimental groups are shown in Figure 3. As expected, HFDPC_K4DT_AR_DHT_ wasabi showed high expression (red color at the bottom of the panel). The names of the 487 genes are listed at the end of Figure 3. In Figure 4, we list the expression profiles of the 344 genes extracted from pairwise comparisons between HFDPC_K4DT_AR_DHT and HFDPC_K4DT_AR_DHT_wasabi. As expected, HFDPC_K4DT_AR_DHT_wasabi showed low expression (blue color at the bottom of the panel). The names of the 344 genes are listed at the end of Figure 4.

**Figure 3.**
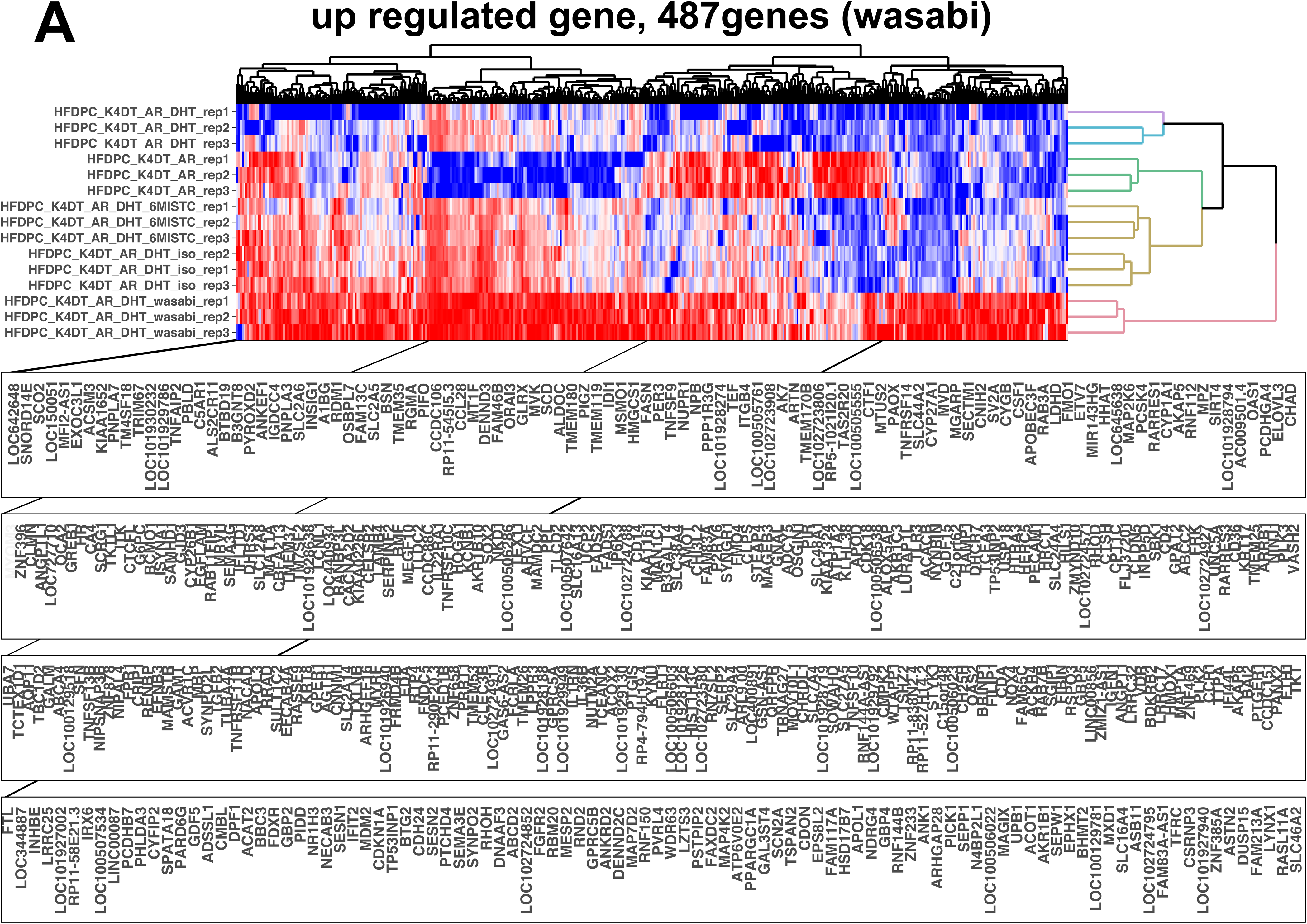
The heatmap analysis of differentially expressed genes (DE genes) of up regulated genes in DHT treated DPCs expressing AR under the exitance of wasabi leaf extract (K4DT_AR_DHT_wasabi). The red indicates the up regulated genes. The blue indicates the down regulated genes.

**Figure 4.**
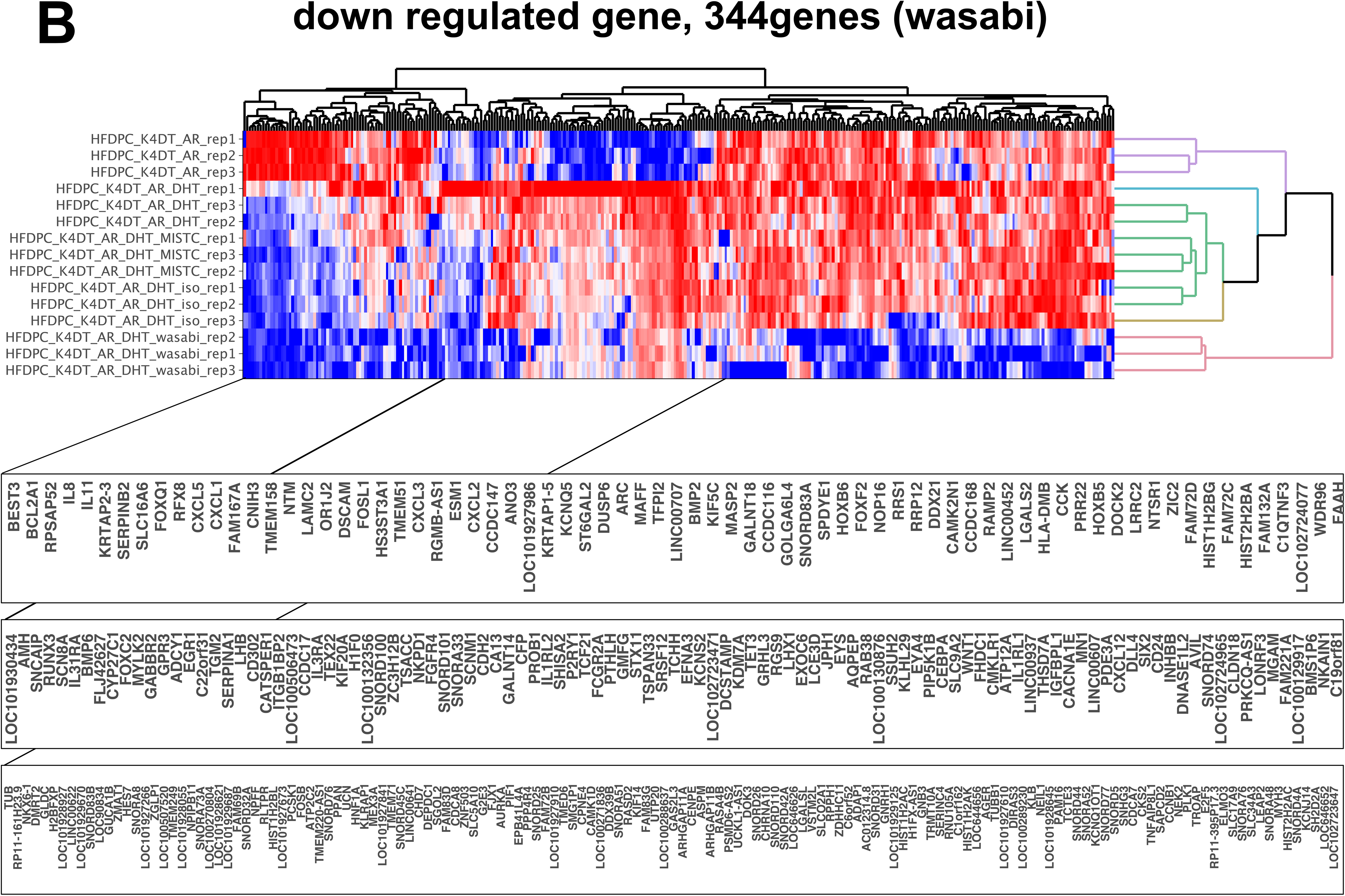
The heatmap analysis of DE genes of down regulated genes in DHT treated DPCs expressing AR under the exitance of wasabi leaf extract (K4DT_AR_DHT_wasabi). The red indicates the up regulated genes. The blue indicates the down regulated genes.

Figure 5 shows a detailed analysis of the DE genes in the isosaponarin-treated group. As shown in Figure 5A, the expression profiles of the 201 upregulated genes were extracted from pairwise comparisons between HFDPC_K4DT_AR_DHT and HFDPC_K4DT_AR_DHT_iso. As expected, the isosaponarin-treated group showed high expression (red color, 7-9^th^ lines from the top). As shown in Figure 5B, the expression profiles of the 148 downregulated genes were extracted from pairwise comparisons. As expected, the isosaponarin-treated group showed low expression (blue color, 4-6^th^ lines from the top). However, the difference in gene expression in isosaponarin was not obvious when compared to the wasabi leaf extract, which could be explained by the small distance of isosaponarin from the HFDPC_K4DT_AR_DHT group in the PCA.

**Figure 5.**
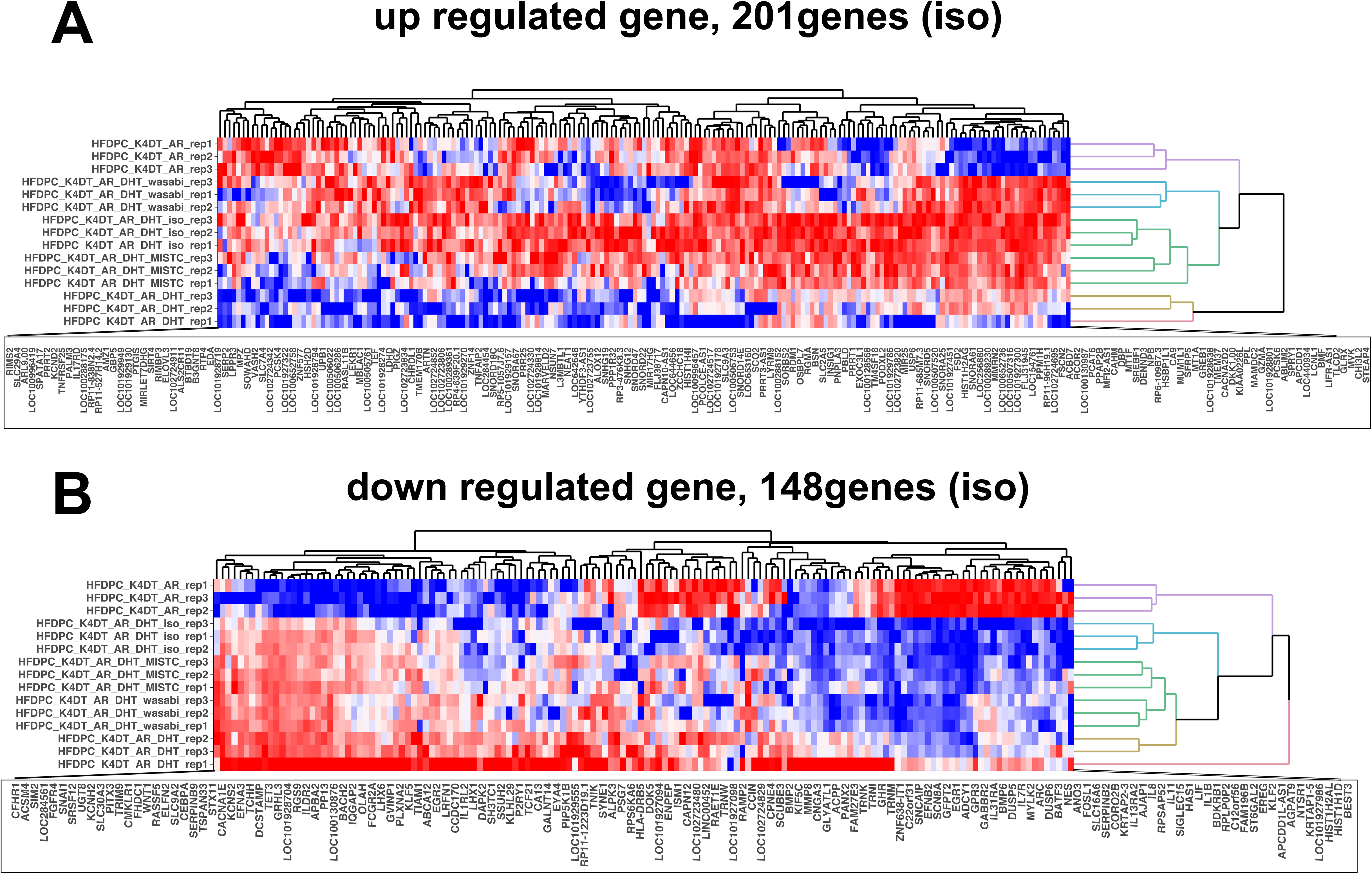
The heatmap analysis of DE genes of up regulated genes in DHT treated DPCs expressing AR under the exitance of isosaponarin. A, Up regulated genes. B, Down regulated genes. The red indicates the up regulated genes. The blue indicates the down regulated genes.

As shown in Figure 6, we performed a detailed analysis of DE genes in the 6-MSITC treated group. The expression profiles of 73 genes were extracted from a pairwise comparison between HFDPC_K4DT_AR_DHT and HFDPC_K4DT_AR_DHT_6-MSITC. As expected, the 6-MSITC treated group showed high expression (red color, bottom line of the panel). The names of the 73 genes are shown in Figure 6A. Figure 6B shows the expression profiles of 67 downregulated genes. As shown in Figure 6B (blue color, 10^th^ line at the top of the panel), 67 genes in the 6-MSITC treated group showed relatively lower expression levels than those in the other experimental groups.

**Figure 6.**
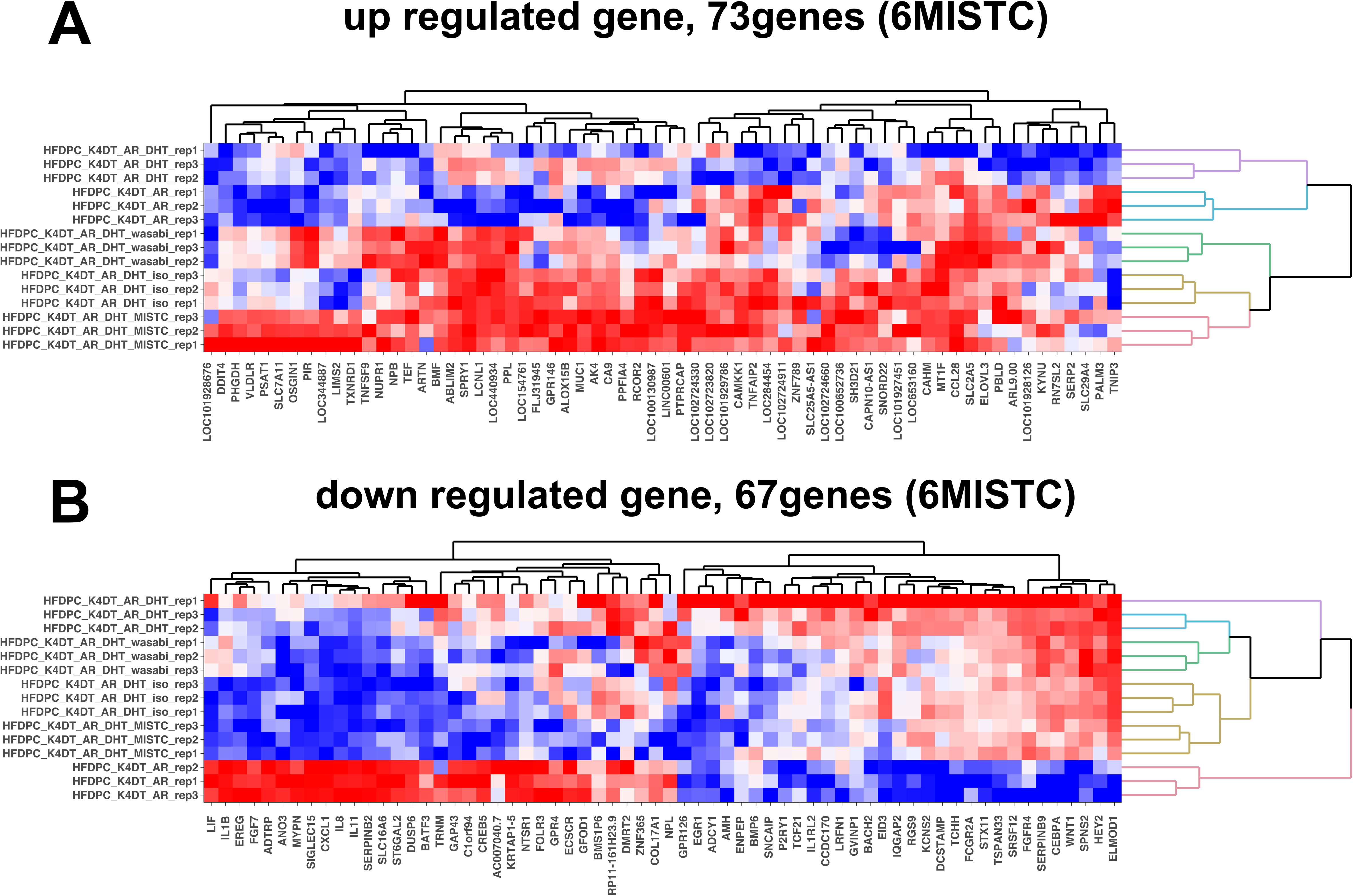
The heatmap analysis of DE genes of up regulated genes in DHT treated DPCs expressing AR under the exitance of 6-MSITC. A, Up regulated genes. B, Down regulated genes. The red indicates the up regulated genes. The blue indicates the down regulated genes.

### Results of the downstream analysis

We conducted a pathway analysis using DAVID based on the DE genes listed earlier. The names of the pathways detected from the upregulated genes in the wasabi-, 6-MSITC-, and isosaponarin-treated groups are listed in Table 1. Furthermore, we listed the names of the pathways detected from the downregulated genes in the wasabi-, 6-MSITC-, and isosaponarin-treated groups in Table 2. The most common characteristics among upregulated and downregulated genes is cytokine-cytokine receptor interaction. In these situations, we highlighted upregulated or downregulated DE genes with arrows derived from the wasabi group within the cytokine-cytokine receptor interaction pathway (Figure 7). Upper arrows indicate upregulated genes. Lower arrows indicate downregulated genes. Detailed information about the expression levels in the cytokine-cytokine receptor interaction pathway are summarized in the bar plots in Figure S1–S13. In the case of increased DE genes, the elevated expression of GDF15 and Decoy receptor 1 (DcR1) was observed in the wasabi-treated group. The role of GDF15 in energy turnover during metabolism is well characterized (Starling, 2023). Transgenic mice expressing GDF15 in fat tissue showed a lower response to a high-fat diet and high sensitivity to insulin (Wang et al., 2023). Furthermore, DcR1 is one of the receptors for the TRAIL ligand, and the activation of TRAIL ligand-receptor signaling results in apoptosis and cell death (Jong et al., 2022). We also observed lower expression levels of bone morphogenetic proteins (BMPs)2. BMP2 play a critical role in hair follicle development or anagen to telogen transition (Botchkarev & Sharov, 2004; Plikus et al., 2008; Wu et al., 2019). The process through which wasabi extract causes high expression of those molecules is not yet known, though there is a possibility that these molecules might be the downstream of the stimulation with wasabi leaf extract.

**Figure 7.**
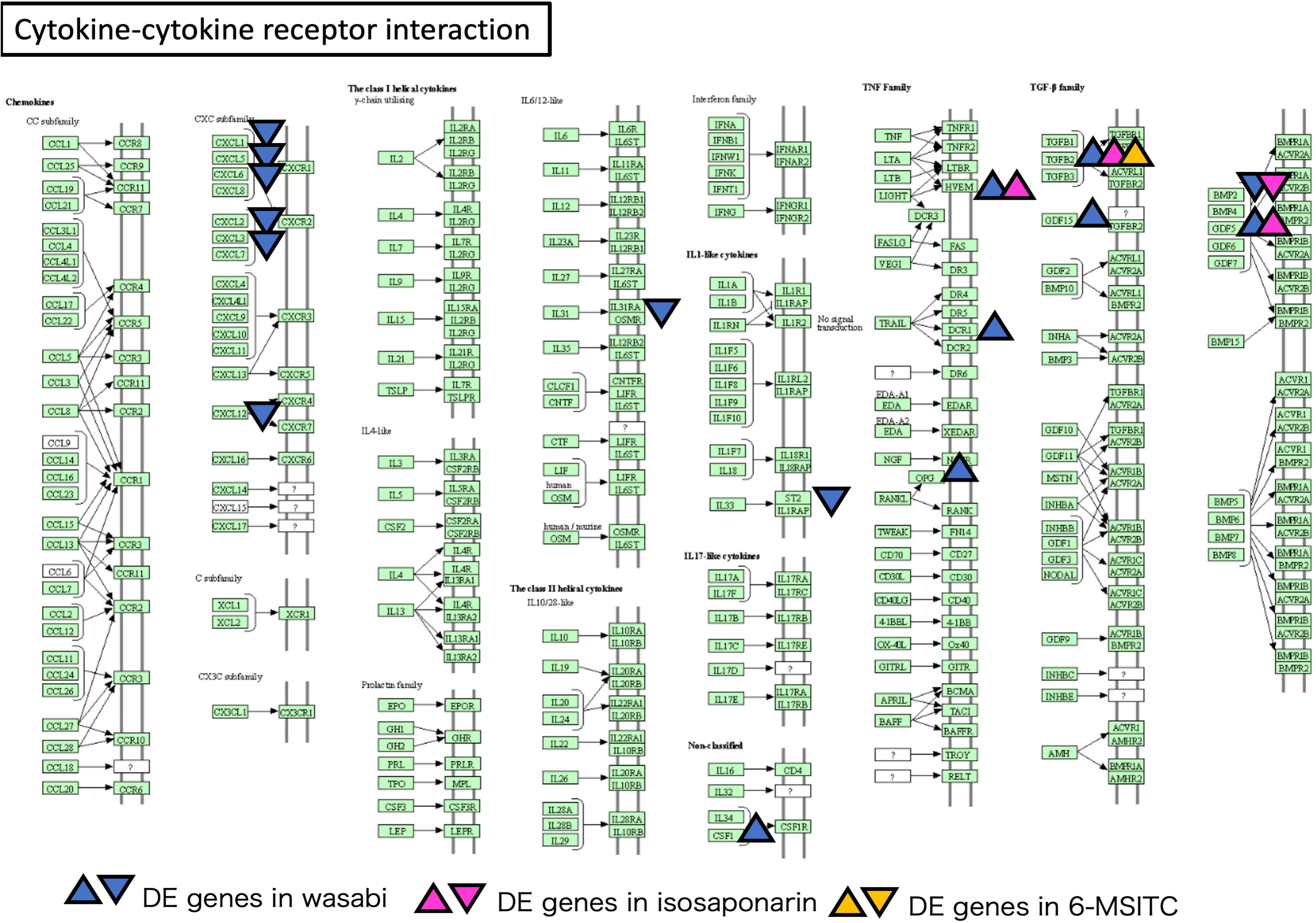
Pathway analysis of Cytokine-cytokine receptor interaction using DAVID and KEGG. The DE genes in wasabi and DE genes in isosaponarin, DE genes in 6-MSITC treated group were shown with upper or lower arrowhead. The direction of the arrows indicate the up regulated or down regulated genes.

Among the down-regulated genes, BMP6 was detected in the wasabi leaf extract-treated group. The role of BMP6 in inhibiting hair follicle stem cell activation has been previously reported. The balance between BMP6 and Wnt10b regulates the telogen-anagen transition in hair follicles (Wu et al., 2019). In addition to BMP6, the downregulation of CXCL1, 2, 3, 5, 6, and 12 was detected in the wasabi leaf-treated group (Figure 7). These ligands belong to the CXC chemokine family (Murdoch & Finn, 2000). CXC chemokines play critical roles in the progression of inflammation (Bikfalvi & Billottet, 2020). CXCL1 is upregulated by the stimulation of inflammation related cytokine, such as IL-1beta (Lee et al., 2015). Therefore, the downregulation of CXC chemokines suggests an anti-inflammatory effect of wasabi leaf extract.

### Chemical analysis of wasabi leaf extract

To detect the polyphenols in the wasabi leaf extract, we performed a chemical analysis. The LC chromatogram of the wasabi leaf extract is shown in Figure 8. Hosoya et al. reported that flavonoids were isolated and identified from wasabi leaves (Hosoya et al., 2005). Suzuki et al. also reported that wasabi contains three types of flavonoids: isosaopnarin, isoorientin, and isovitexin, which are mainly found in its leaves and flowers (Mashima et al., 2019). In particular, wasabi leaves contain high concentrations of isosaponarin and phenolic acids (Szewczyk et al., 2021). In this study, we attempted to identify three major peaks in the chromatogram. The chemical formulas of isosaponalin, isoorientin, and isovitexin are C27H30O15, C21H20O11, and C21H20O10, respectively. Therefore, the protonated molecular ions [M+H]^+^ were calculated as 595.1657 (isosaponalin), 449.1078 (isoorientin), and 433.1129 (isovitexin) from each monoisotopic mass. The peaks corresponding to the retention times of the LC chromatogram were obtained from the extracted ion chromatograms of [M+H]^+^ (Figure 8). Furthermore, isosaponarin, isovitexin, and isoorientin were identified at each retention time using standard substances. In addition to these flavonoids, other main peaks have not yet been identified.

**Figure 8.**
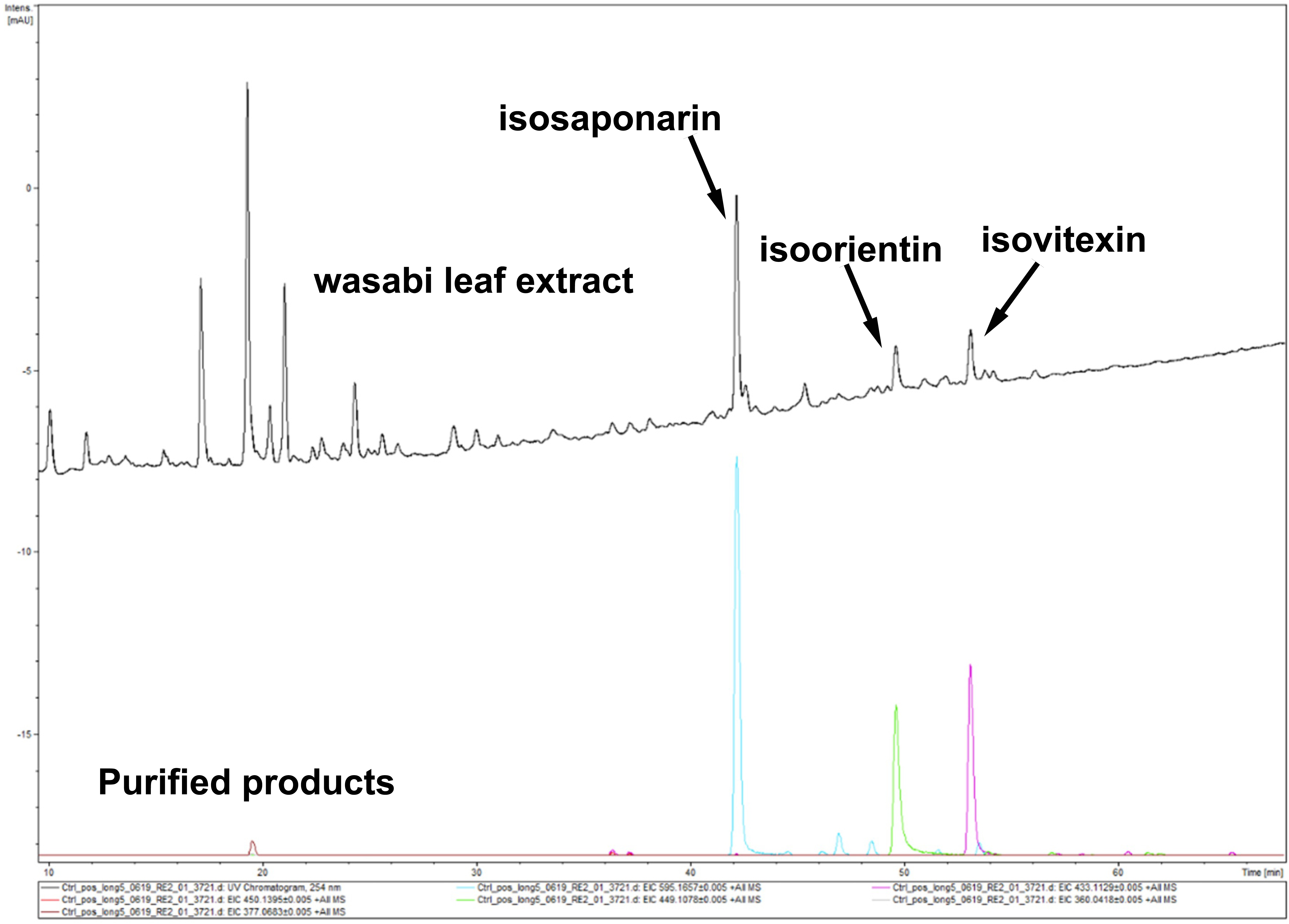
Chemical analysis of wasabi leaf extract used for this study. HPLC chart or purified products. The peaks which possibly derived from isosaponarin, isoorientin, isovitexin were indicated by arrows.

## DATA AVAILABILITY

The data that support the findings of this study are available from the corresponding author, [T.F.], upon reasonable request.

## Ethic statement and IRB statement

In this study, we used immortalized DPCs from human. However, the original primary DPCs were commercially obtained. Based on the judgement of committee of iwate university, approval of human related materials is not required for this study.

## ACKNOWLEDGEMENTS

We would like to thank Professor Goro Taguchi (Sinshu University, Faculty of Textile Science and Technology, Department of Applied Biology) for providing technical help during the purification of isosaponarin.

## AUTHOR CONTRIBUTION

HM, TYK, MN, LB, TF contributed to the data collection. HM, TYK, MN, LB, TF did the data analysis. HM, TYK, HT, ES, TO, IO, TF contributed to the study design. HM, TYK, IO, TF prepared the manuscript.

## LEGENDS FOR THE SUPPLIMENTAL MATERIALS

**Figure S1.**
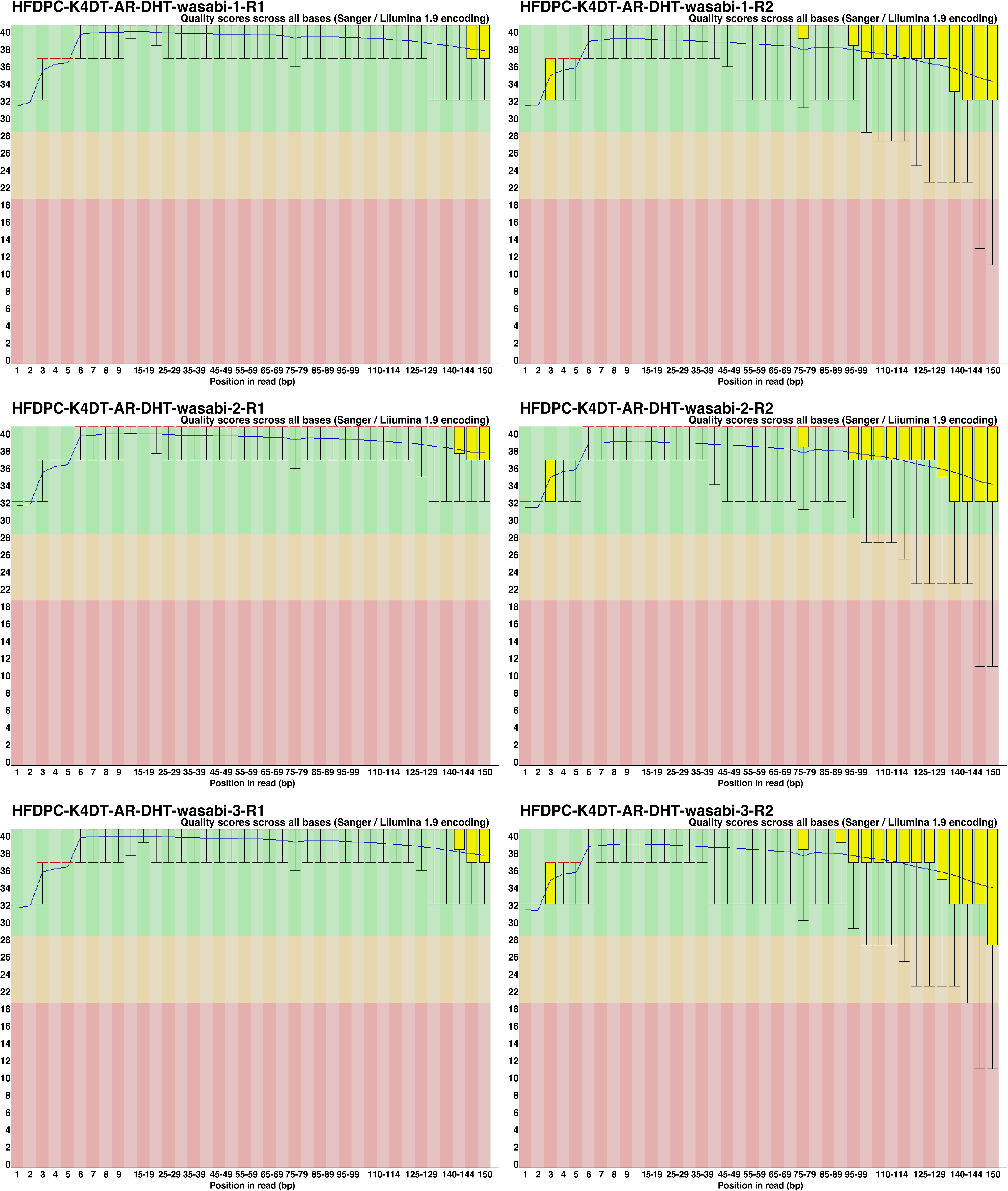
Results of read quality validation using FASTQC in wasabi leaf extract treated group.

**Figure S2.**
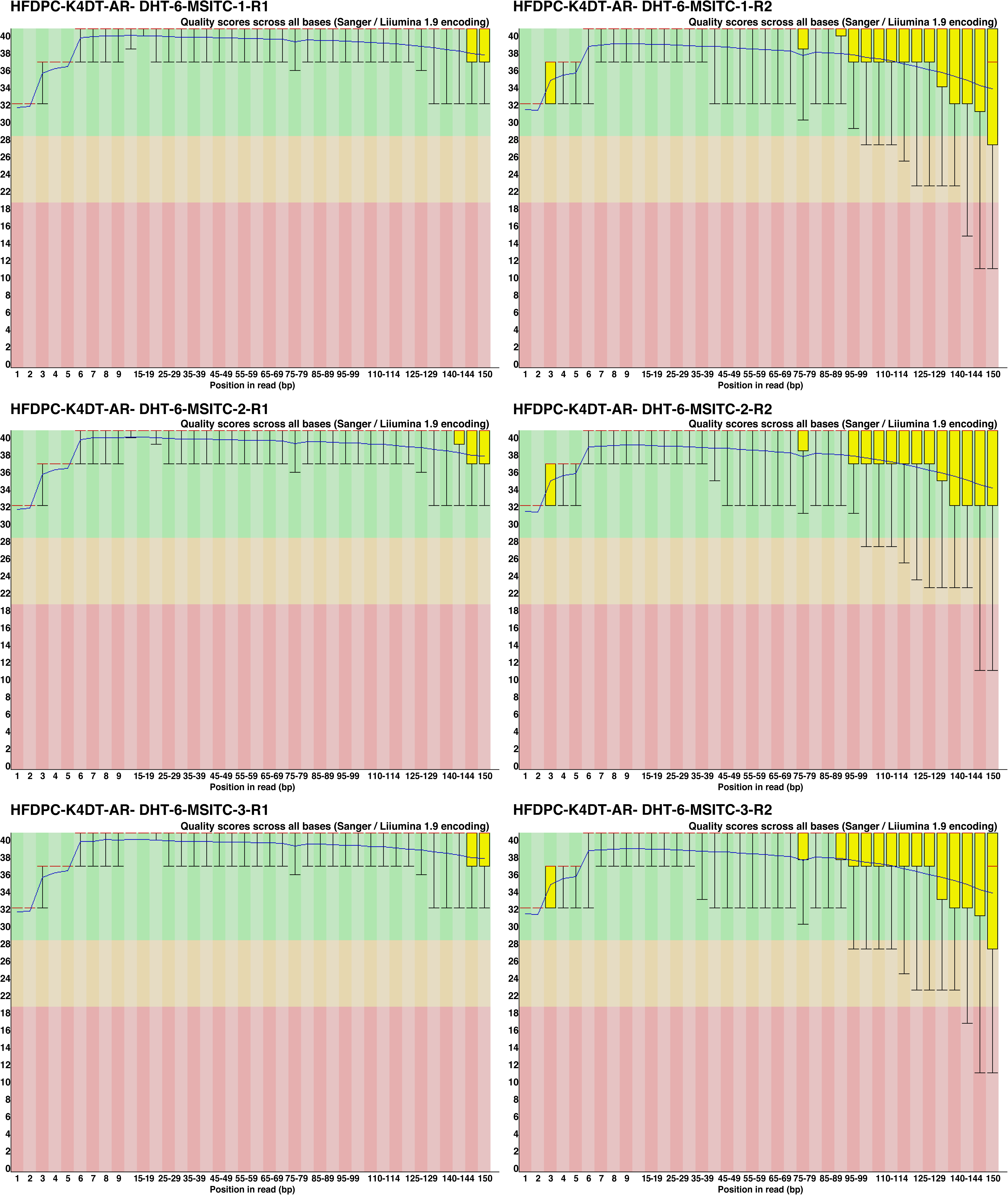
Results of read quality validation using FASTQC in 6-MSITC treated group.

**Figure S3.**
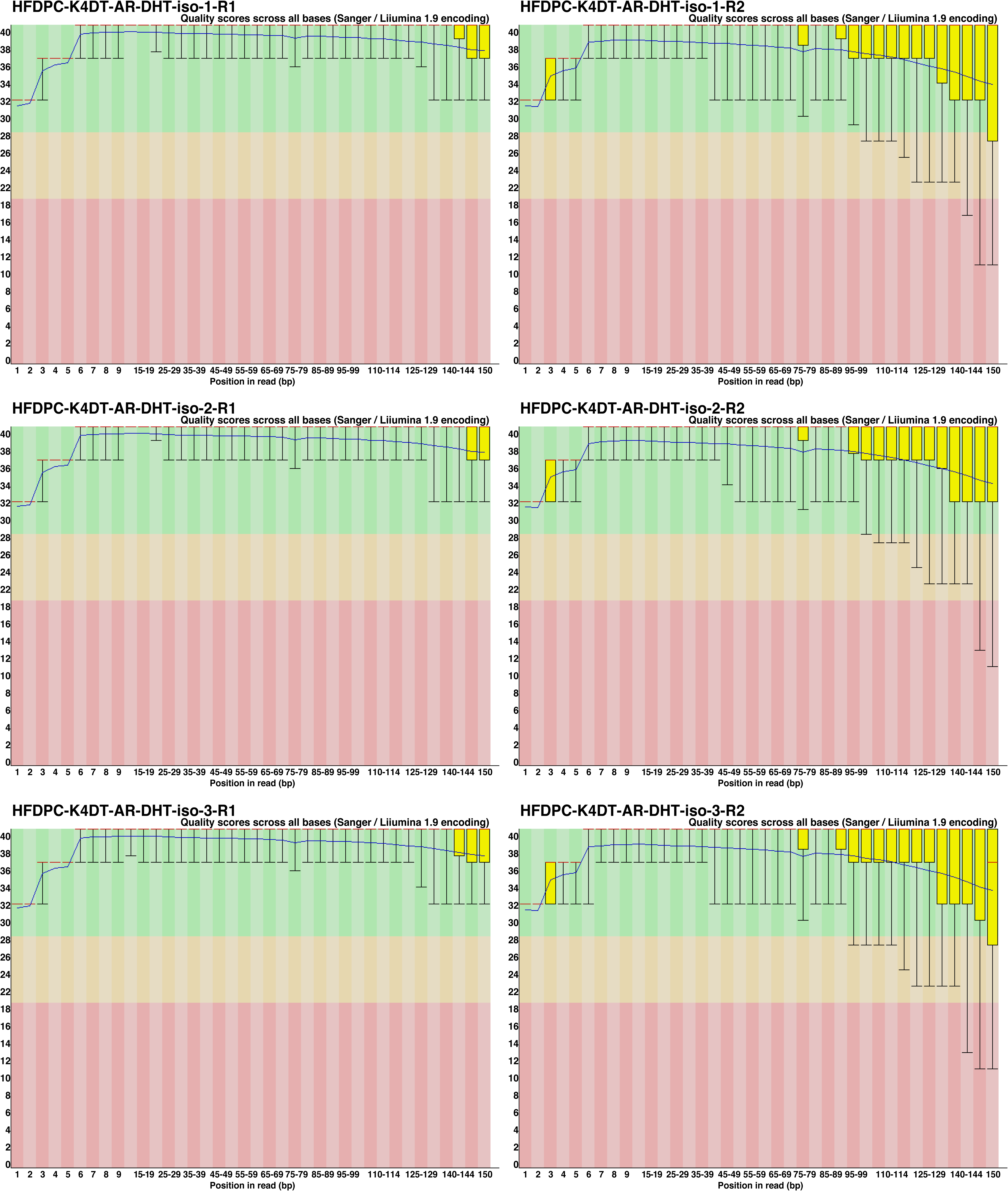
Results of read quality validation using FASTQC in isosaponarin treated group.

**Figure S4.**
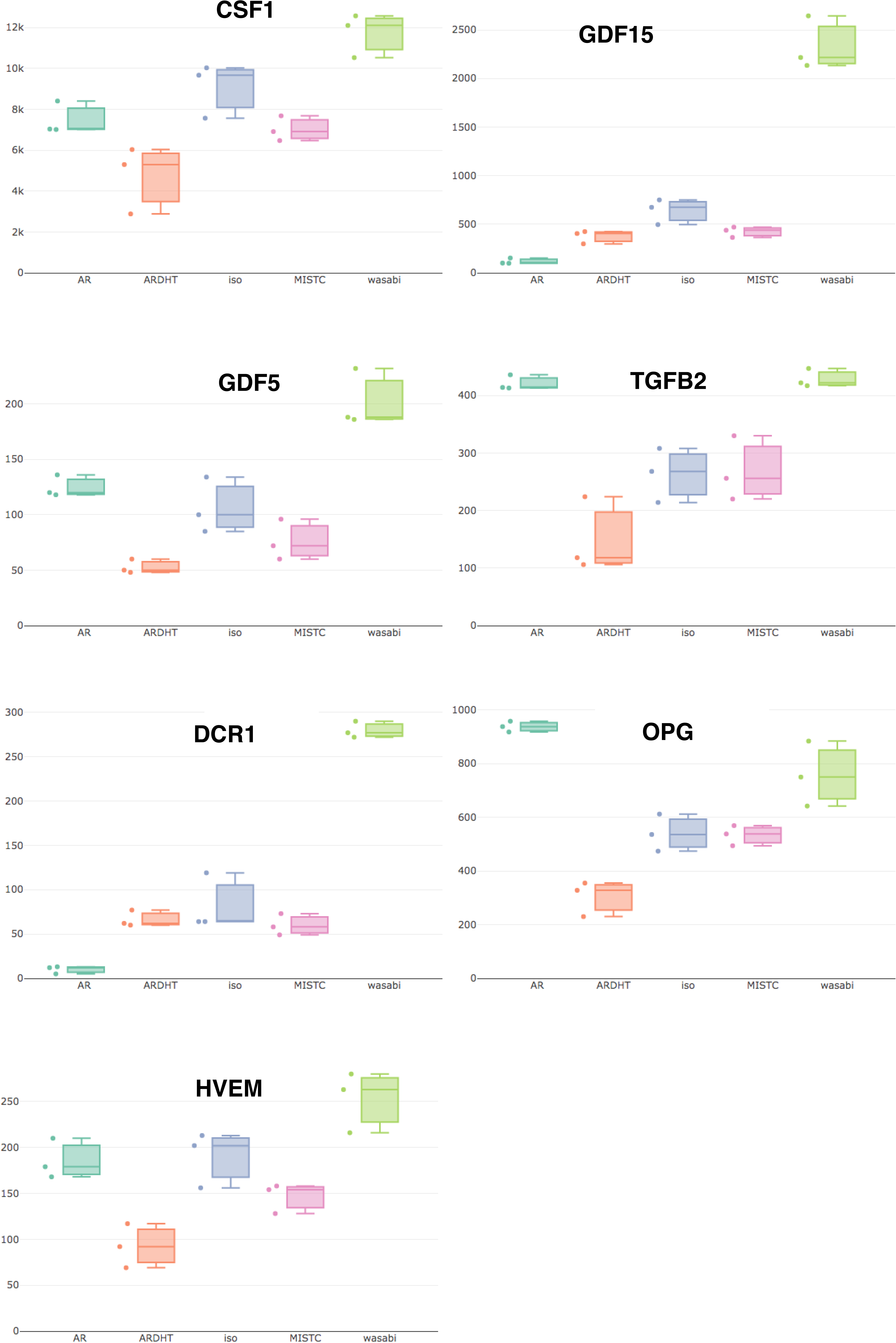
Bar plots of upregulated genes in wasabi group on Cytokine-cytokine receptor interaction pathway.

**Figure S5.**
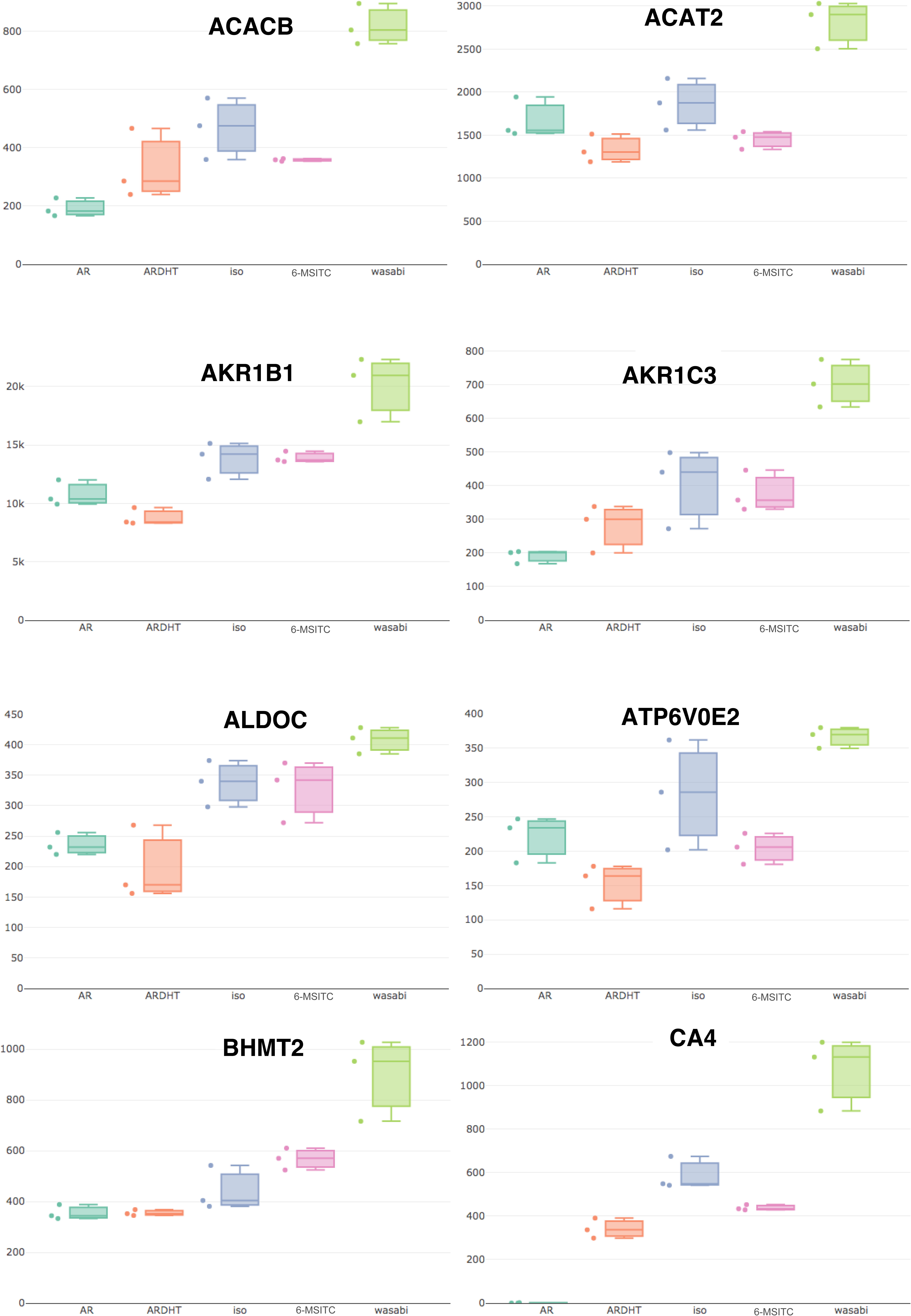
Bar plots of upregulated genes in wasabi group on Metabolic Pathway. Part 1.

**Figure S6.**
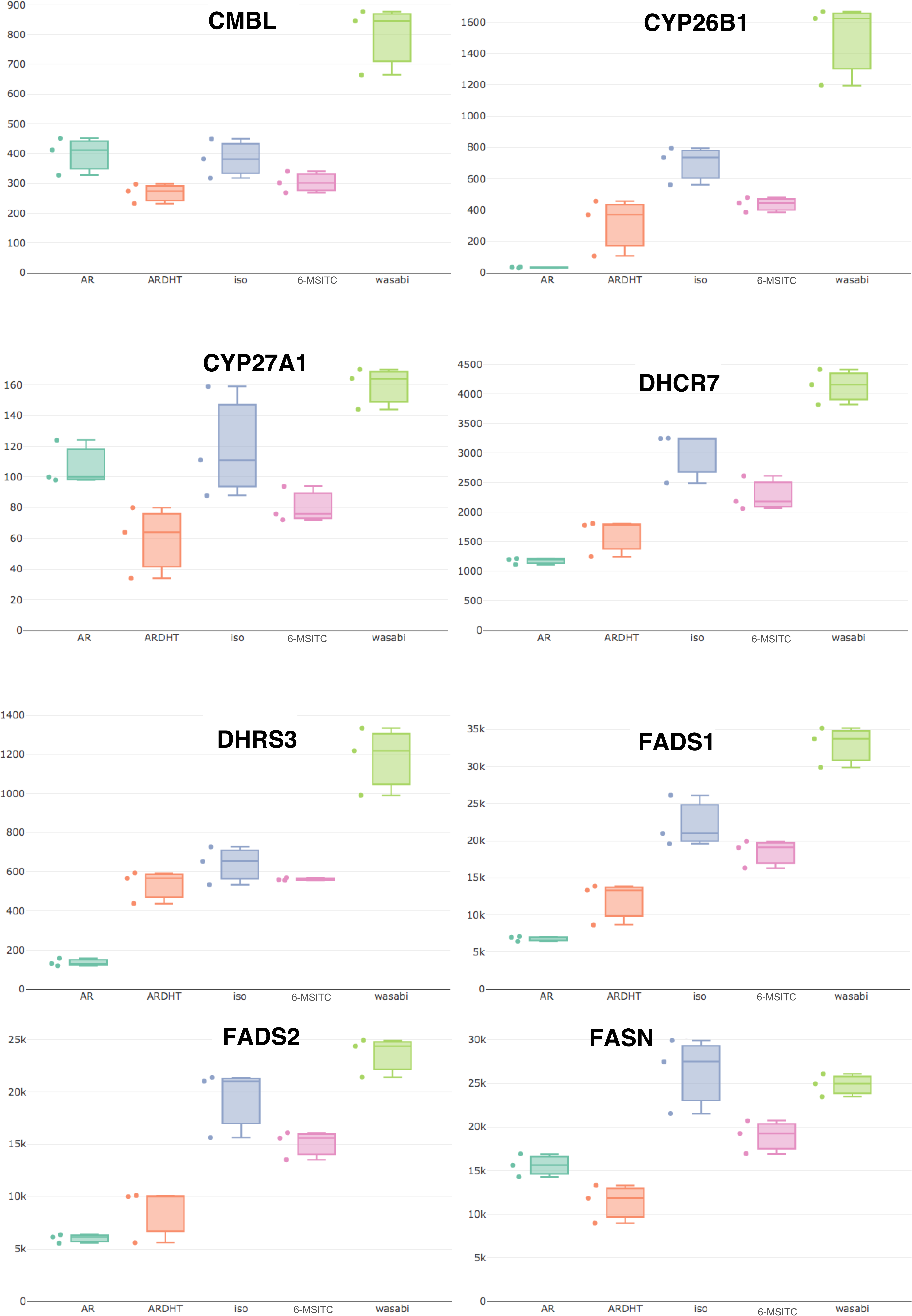
Bar plots of upregulated genes in wasabi group on Metabolic Pathway. Part 2.

**Figure S7.**
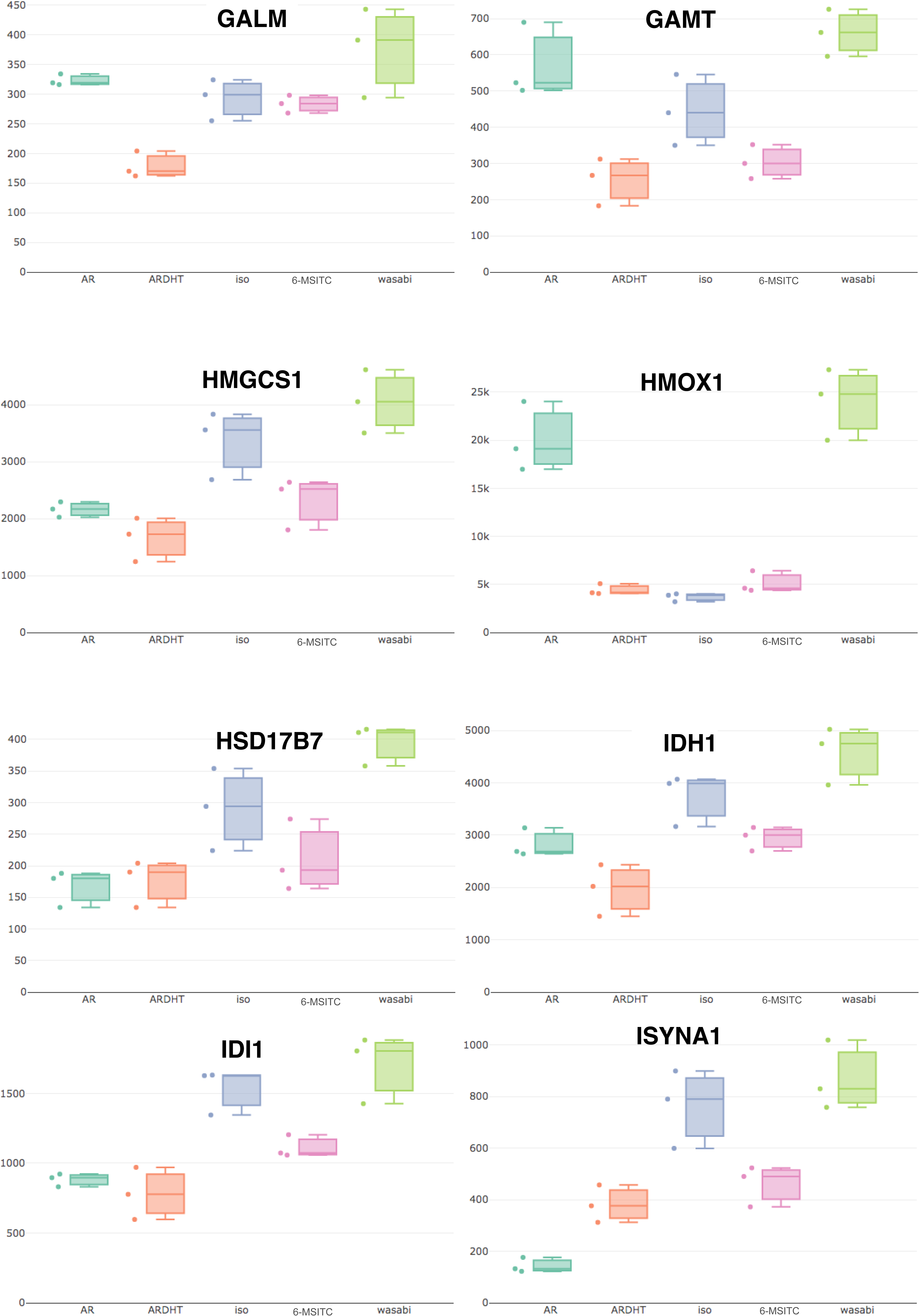
Bar plots of upregulated genes in wasabi group on Metabolic Pathway. Part 3.

**Figure S8.**
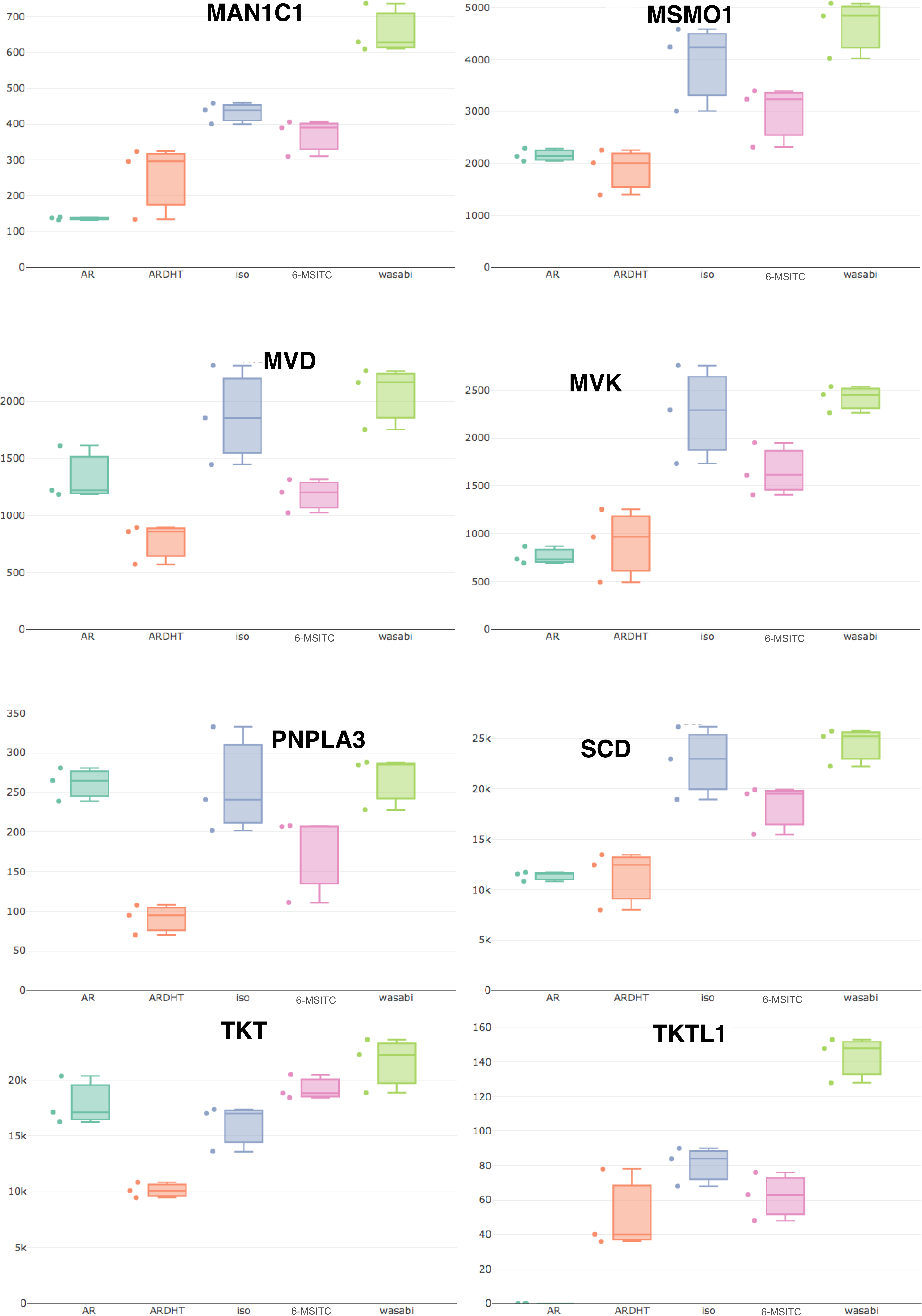
Bar plots of upregulated genes in wasabi group on Metabolic Pathway. Part 4.

**Figure S9.**
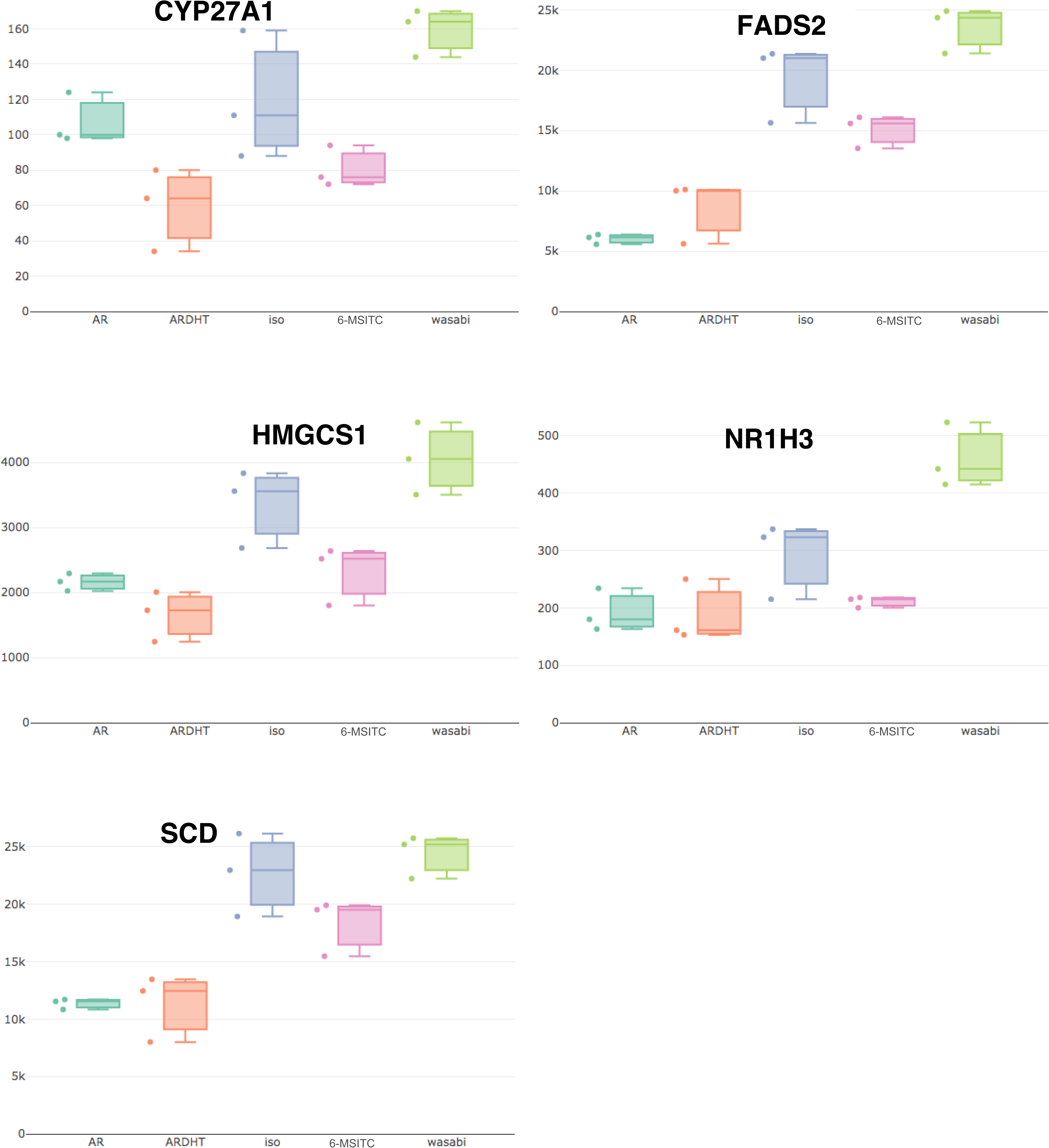
Bar plots of upregulated genes in wasabi group on PPAR signaling pathway.

**Figure S10.**
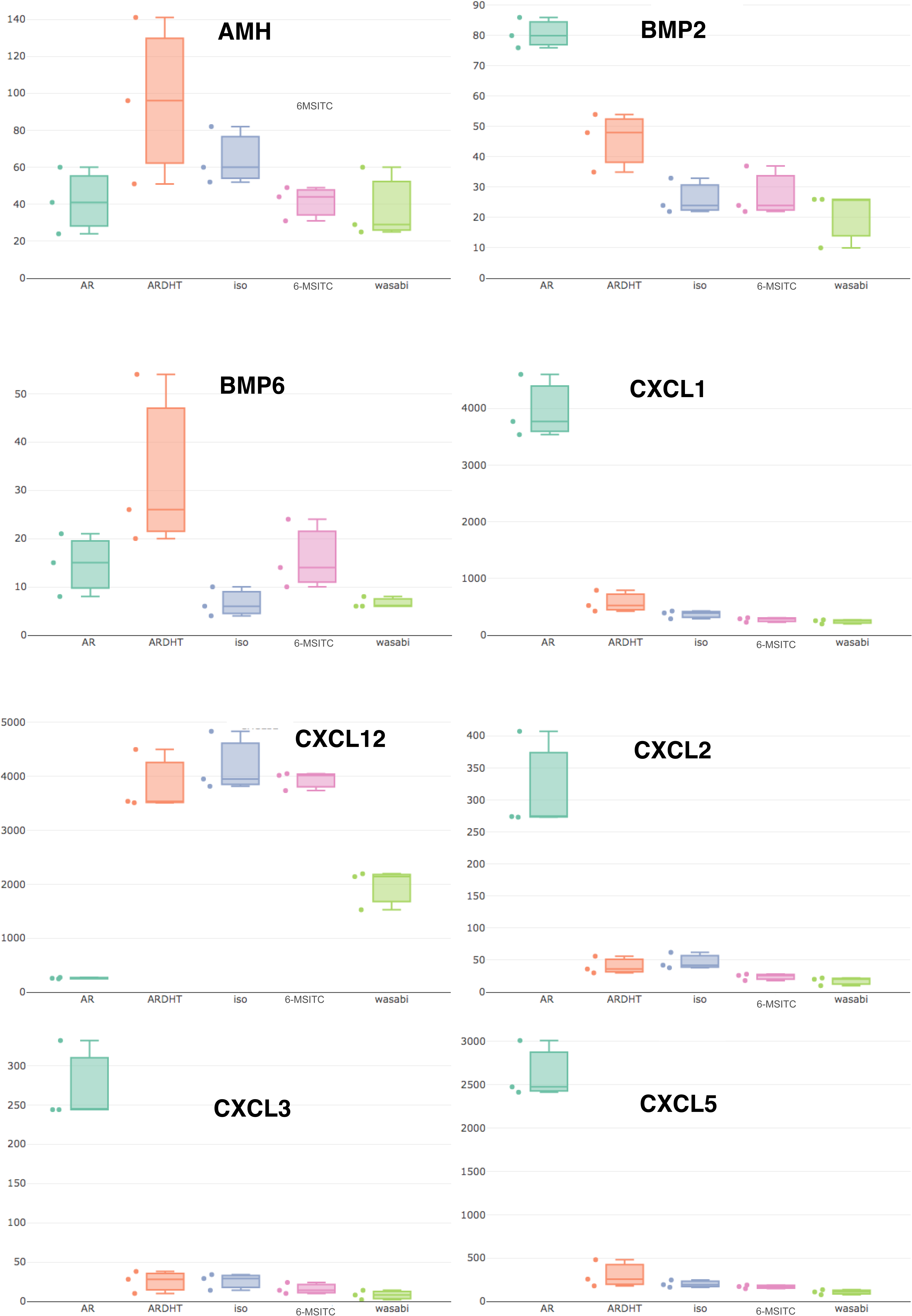
Bar plots of downregulated genes in wasabi group on Cytokine-cytokine receptor interaction pathway. Part 1.

**Figure S11.**
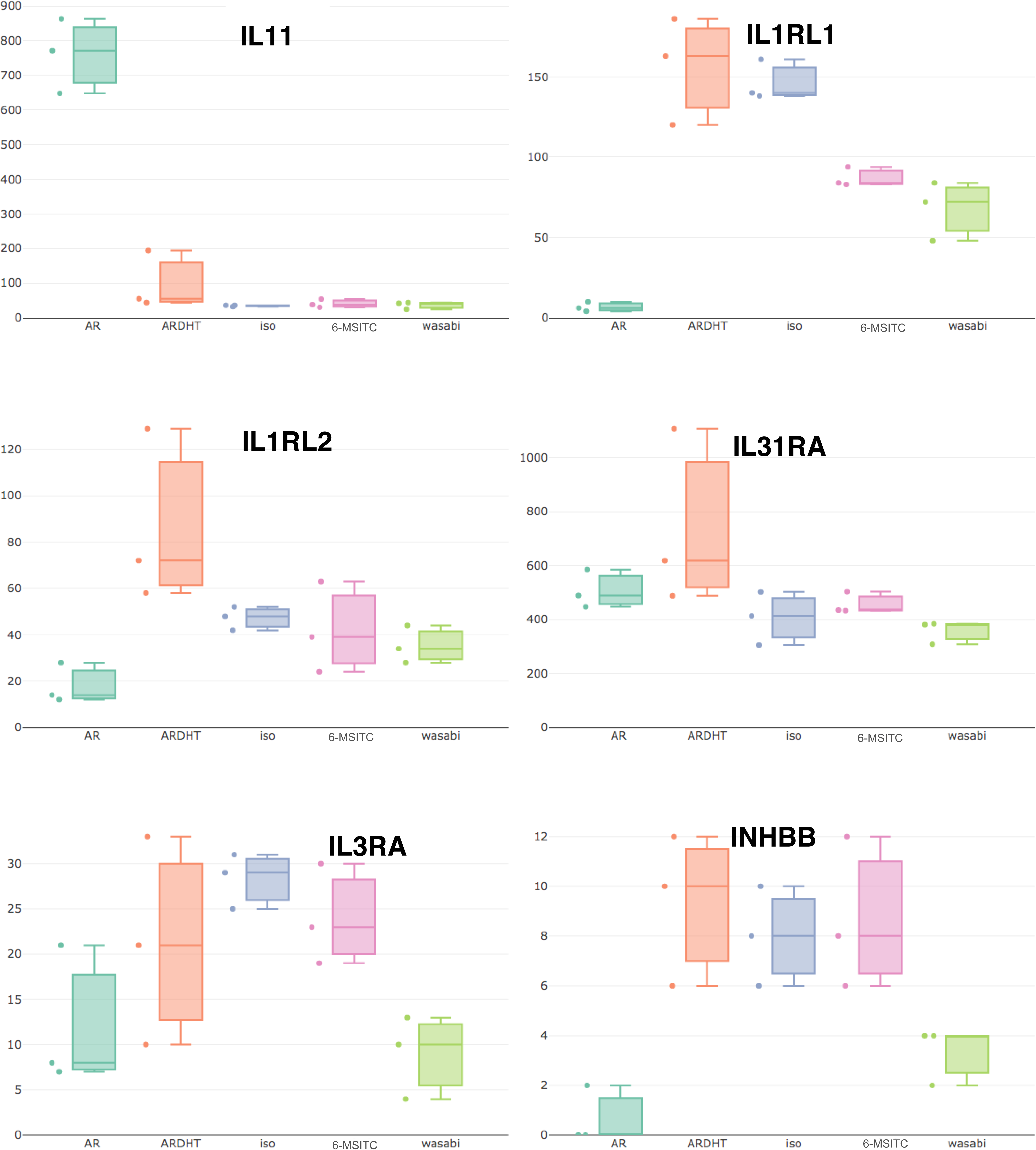
Bar plots of downregulated genes in wasabi group on Cytokine-cytokine receptor interaction pathway. Part 2.

**Figure S12.**
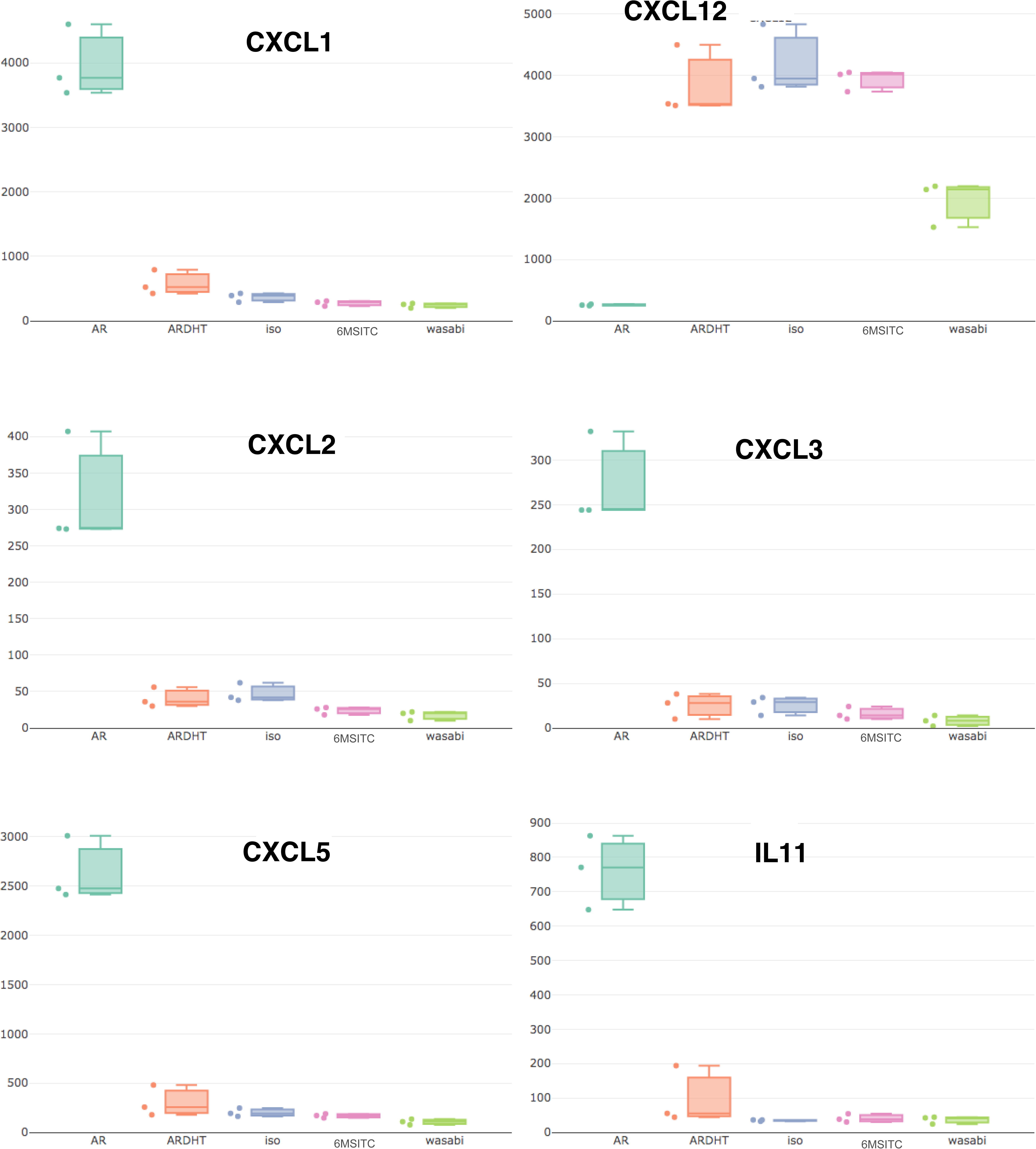
Bar plots of downregulated genes in wasabi group on Rheumatoid arthritis pathway.

**Figure S13.**
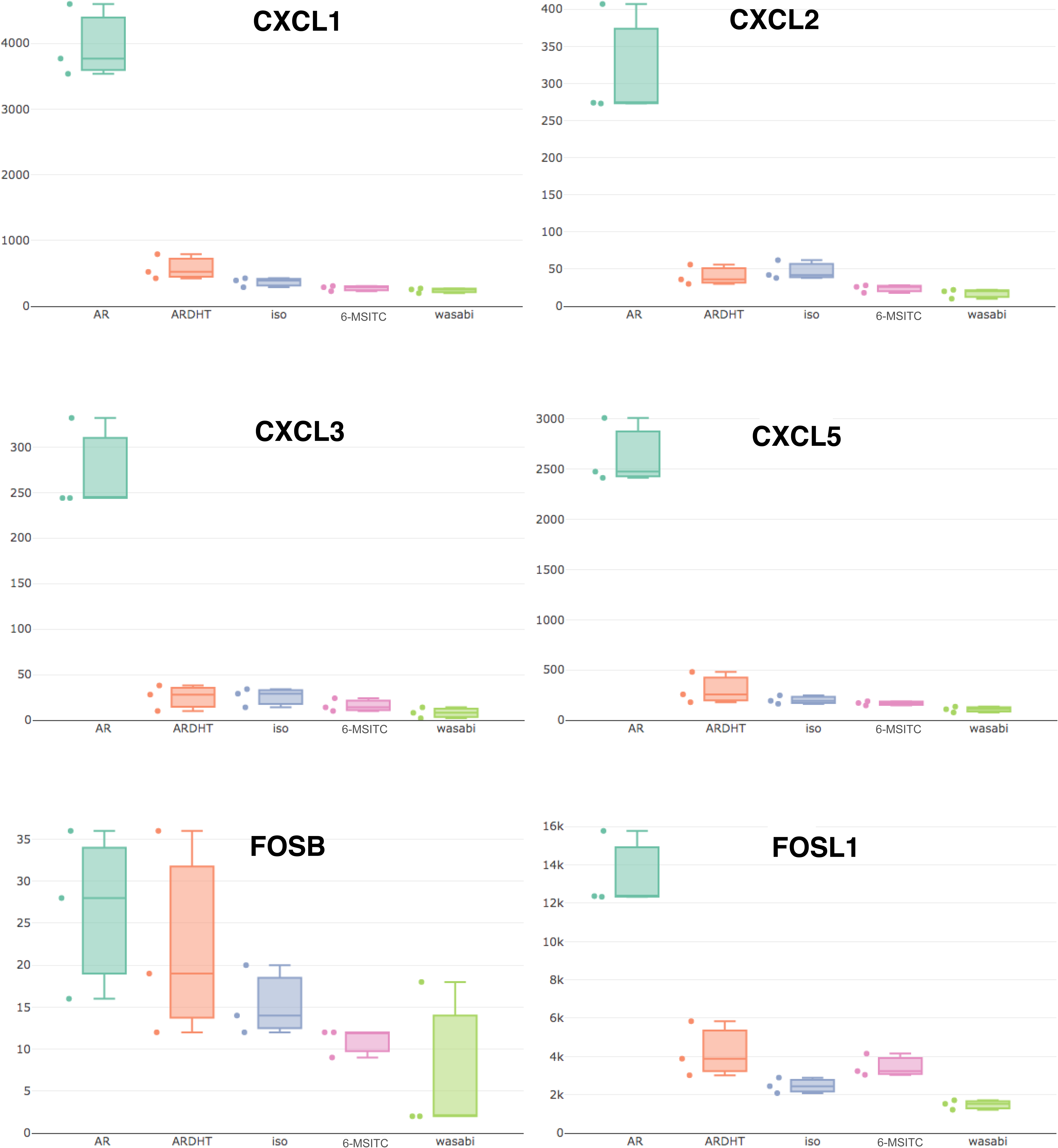
Bar plots of downregulated genes in wasabi group on IL-17 signaling pathway.

## REFERENCE

Bikfalvi, A., & Billottet, C. (2020). Cellular Processes in Tumor Metastasis: From Basic Research to Translation: The CC and CXC chemokines: major regulators of tumor progression and the tumor microenvironment. American Journal of Physiology - Cell Physiology, 318(3), C542. 10.1152/AJPCELL.00378.2019

Botchkarev, V. A., & Sharov, A. A. (2004). BMP signaling in the control of skin development and hair follicle growth. Differentiation, 72(9–10), 512–526. 10.1111/J.1432-0436.2004.07209005.X

Chen, S., Zhou, Y., Chen, Y., & Gu, J. (2018). fastp: an ultra-fast all-in-one FASTQ preprocessor. Bioinformatics, 34(17), i884–i890. 10.1093/BIOINFORMATICS/BTY560

Dobin, A., Davis, C. A., Schlesinger, F., Drenkow, J., Zaleski, C., Jha, S., Batut, P., Chaisson, M., & Gingeras, T. R. (2013). STAR: ultrafast universal RNA-seq aligner. Bioinformatics, 29(1), 15–21. 10.1093/BIOINFORMATICS/BTS635

Fukuda, T., Furuya, K., Takahashi, K., Orimoto, A., Sugano, E., Tomita, H., Kashiwagi, S., Kiyono, T., & Ishii, T. (2021). Combinatorial expression of cell cycle regulators is more suitable for immortalization than oncogenic methods in dermal papilla cells. IScience, 24(1), 101929. 10.1016/j.isci.2020.101929

Fukuda, T., Takahashi, K., Takase, S., Orimoto, A., Eitsuka, T., Nakagawa, K., & Kiyono, T. (2020). Human Derived Immortalized Dermal Papilla Cells With a Constant Expression of Testosterone Receptor. Frontiers in Cell and Developmental Biology, 8. 10.3389/FCELL.2020.00157

Furuya, K., Fujibayashi, S., Wu, T., Takahashi, K., Takase, S., Orimoto, A., Sugano, E., Tomita, H., Kashiwagi, S., Kiyono, T., Ishii, T., & Fukuda, T. (2022). Transcriptome analysis to identify the downstream genes of androgen receptor in dermal papilla cells. BMC Genomic Data, 23(1). 10.1186/S12863-021-01018-6

Hosoya, T., Young, S. Y., & Kunugi, A. (2005). Five novel flavonoids from Wasabia japonica. Tetrahedron, 61(29), 7037–7044. 10.1016/J.TET.2005.04.061

Jong, K. X. J., Mohamed, E. H. M., & Ibrahim, Z. A. (2022). Escaping cell death via TRAIL decoy receptors: a systematic review of their roles and expressions in colorectal cancer. Apoptosis : An International Journal on Programmed Cell Death, 27(11–12), 787–799. 10.1007/S10495-022-01774-5

Lee, Y. S., Yang, H., Yang, J. Y., Kim, Y., Lee, S. H., Kim, J. H., Jang, Y. J., Vallance, B. A., & Kweon, M. N. (2015). Interleukin-1 (IL-1) signaling in intestinal stromal cells controls KC/ CXCL1 secretion, which correlates with recruitment of IL-22- secreting neutrophils at early stages of Citrobacter rodentium infection. Infection and Immunity, 83(8), 3257–3267. 10.1128/IAI.00670-15

Li, Y.-L., Weng, J.-C., Hsiao, C.-C., Chou, M.-T., Tseng, C.-W., & Hung, J.-H. (2015). PEAT: an intelligent and efficient paired-end sequencing adapter trimming algorithm. BMC Bioinformatics, 16 Suppl 1(Suppl 1), S2. 10.1186/1471-2105-16-S1-S2

Liao, Y., Smyth, G. K., & Shi, W. (2014). featureCounts: an efficient general purpose program for assigning sequence reads to genomic features. Bioinformatics, 30(7), 923–930. 10.1093/BIOINFORMATICS/BTT656

Mashima, K., Hatano, M., Suzuki, H., Shimosaka, M., & Taguchi, G. (2019). Identification and Characterization of Apigenin 6-C-Glucosyltransferase Involved in Biosynthesis of Isosaponarin in Wasabi (Eutrema japonicum). Plant & Cell Physiology, 60(12), 2733–2743. 10.1093/PCP/PCZ164

Matsusaka, H., Wu, T., Furuya, K., Yamada-Kato, T., Bai, L., Tomita, H., Sugano, E., Ozaki, T., Kiyono, T., Okunishi, I., & Fukuda, T. (2022). Comprehensive transcriptome data to identify downstream genes of testosterone signalling in dermal papilla cells. Scientific Data, 9(1). 10.1038/S41597-022-01846-W

Morgan, L. (2005). Growing Edge International. https://books.google.co.jp/books?id=lZD95wlLhxIC&pg=PA53&redir_esc=y#v=onepage&q&f=false

Murdoch, C., & Finn, A. (2000). Chemokine receptors and their role in inflammation and infectious diseases. Blood, 95(10), 3032–3043. 10.1182/BLOOD.V95.10.3032

Nagai, M., Akita, K., Yamada, K., & Okunishi, I. (2010). The effect of isosaponarin isolated from wasabi leaf on collagen synthesis in human fibroblasts and its underlying mechanism. Journal of Natural Medicines, 64(3), 305–312. 10.1007/S11418-010-0412-Y/FIGURES/13

Orimoto, A., Kyakumoto, S., Eitsuka, T., Nakagawa, K., Kiyono, T., & Fukuda, T. (2020). Efficient immortalization of human dental pulp stem cells with expression of cell cycle regulators with the intact chromosomal condition. PloS One, 15(3). 10.1371/JOURNAL.PONE.0229996

Orimoto, A., Takahashi, K., Imai, M., Kiyono, T., Kawaoka, Y., & Fukuda, T. (2021). Establishment of human airway epithelial cells with doxycycline-inducible cell growth and fluorescence reporters. Cytotechnology, 73(4), 555–569. 10.1007/S10616-021-00477-0

Plikus, M. V., Mayer, J. A., De La Cruz, D., Baker, R. E., Maini, P. K., Maxson, R., & Chuong, C. M. (2008). Cyclic dermal BMP signalling regulates stem cell activation during hair regeneration. Nature 2008 451:7176, 451(7176), 340–344. 10.1038/nature06457

Shiomi, K., Kiyono, T., Okamura, K., Uezumi, M., Goto, Y., Yasumoto, S., Shimizu, S., & Hashimoto, N. (2011). CDK4 and cyclin D1 allow human myogenic cells to recapture growth property without compromising differentiation potential. Gene Therapy, 18(9), 857–866. 10.1038/GT.2011.44

Shizuoka. (2023). Traditional Cultivation of Shizuoka Water Wasabi” was recognized as a World Agricultural Heritage Site! | Shizuoka Prefecture Official Website. Http://Www_dr.Pref.Shizuoka.Jp/Kurashikankyo/Shokuseikatsu/Kaju/1003330/1027389.Html. http://www_dr.pref.shizuoka.jp/kurashikankyo/shokuseikatsu/kaju/1003330/1027389.html

Starling, S. (2023). GDF15 boosts muscle energy burn. Nature Reviews. Endocrinology, 19(9), 499. 10.1038/S41574-023-00877-6

Su, W., Sun, J., Shimizu, K., & Kadota, K. (2019). TCC-GUI: a Shiny-based application for differential expression analysis of RNA-Seq count data. BMC Research Notes, 12(1), 133. 10.1186/s13104-019-4179-2

Szewczyk, K., Pietrzak, W., Klimek, K., Miazga-Karska, M., Firlej, A., Flisiński, M., & Grzywa-Celińska, A. (2021). Flavonoid and phenolic acids content and in vitro study of the potential anti-aging properties of eutrema japonicum (Miq.) koidz cultivated in wasabi farm poland. International Journal of Molecular Sciences, 22(12), 6219. 10.3390/IJMS22126219/S1

Trio, P. Z., Kawahara, A., Tanigawa, S., Sakao, K., & Hou, D. X. (2017). DNA Microarray Profiling Highlights Nrf2-Mediated Chemoprevention Targeted by Wasabi-Derived Isothiocyanates in HepG2 Cells. Nutrition and Cancer, 69(1), 105–116. 10.1080/01635581.2017.1248296

Wang, D., Townsend, L. K., DesOrmeaux, G. J., Frangos, S. M., Batchuluun, B., Dumont, L., Kuhre, R. E., Ahmadi, E., Hu, S., Rebalka, I. A., Gautam, J., Jabile, M. J. T., Pileggi, C. A., Rehal, S., Desjardins, E. M., Tsakiridis, E. E., Lally, J. S. V., Juracic, E. S., Tupling, A. R., … Steinberg, G. R. (2023). GDF15 promotes weight loss by enhancing energy expenditure in muscle. Nature, 619(7968), 143–150. 10.1038/S41586-023-06249-4

Watanabe, M., Ohata, M., Hayakawa, S., Isemura, M., Kumazawa, S., Nakayama, T., Furugori, M., & Kinae, N. (2003). Identification of 6-methylsulfinylhexyl isothiocyanate as an apoptosis-inducing component in wasabi. Phytochemistry, 62(5), 733–739. 10.1016/S0031-9422(02)00613-1

Wu, P., Zhang, Y., Xing, Y., Xu, W., Guo, H., Deng, F., Ma, X., & Li, Y. (2019). The balance of Bmp6 and Wnt10b regulates the telogen-anagen transition of hair follicles. Cell Communication and Signaling, 17(1), 1–10. 10.1186/S12964-019-0330-X/FIGURES/6

